# Fish Show Genetic Evolutionary Responses to River Regulation

**DOI:** 10.1101/2025.08.01.668107

**Authors:** Xiatong Cai, Io Deflem, Colin D. Rennie, Joke De Meester, Patrick Willems, Andrew P. Hendry, Federico C.F. Calboli, Bart Hellemans, Filip A.M. Volckaert, Joost A.M. Raeymaekers

**Affiliations:** Department of Civil Engineering, University of Ottawa, 161 Louis-Pasteur Private, K1N 6N5, Ottawa, ON, Canada; Faculty of Biosciences and Aquaculture, Nord University, Universitetsalléen 11, N-8026 Bodø, Norway; Research Institute for Nature and Forest (INBO), B-9500 Geraardsbergen, Belgium; Hydraulics and Geotechnics Section, KU Leuven, Kasteelpark Arenberg 40, BE-3001, Leuven, Belgium; Flanders Hydraulics Research, Department of Mobility and Public Works, Flemish Government, Berchemlei 115, 2140 Antwerp, Belgium; Department of Biology, Redpath Museum, McGill University, 859 Sherbrooke Street West, H3A 0C4, Montreal, Quebec, Canada; Natural Resources Institute Finland, Latokartanokaari 9, 00790 Helsinki, Finland; Laboratory of Biodiversity and Evolutionary Genomics, KU Leuven, B-3000 Leuven, Belgium

**Keywords:** eco-hydraulics, fish, evolution, adaptive potential, eco-evo-hydraulics

## Abstract

Eco-hydraulics traditionally aims at managing riverine systems in a semi-natural state while meeting human demands, assuming aquatic species are static. However, evidence of rapid evolution suggests that ignoring evolutionary dynamics of fish species might limit long-term effectiveness of eco-hydraulics frameworks. It remains unclear how freshwater fish adapt to human perturbation. Why are some fish populations more resilient to human perturbation than others? What are the genetic mechanisms behind it? To answer these questions, we genotyped eleven populations of three-spined stickleback in a regulated river system and collected data on river morphology, connectivity, flow regimes, physico-chemistry and parasite abundance through a combination of field surveys and modelling. Gene-environment association analysis detected strong signals of genetic divergence associated with hydraulic features. Gene ontology analysis revealed evolutionary responses that primarily involve functions in the nervous and sensory systems. These findings demonstrate that fish can evolve in response to river regulation, highlighting the need to transition from eco-hydraulics toward eco-evo-hydraulics.

---

Globally, freshwater fish biodiversity is facing a tremendous decline. Recently, a study examining the International Union for Conservation of Nature’s (IUCN) Red List of Threatened Species pointed out that 26% freshwater fishes are threatened with extinction^1^. In Europe, there are at least 1.2 million instream barriers with a mean density of 0.74 barriers per kilometre, of which 68% are less than two meters^2^. These barriers restrict fish movement and enlarge the genetic divergence within water basins^3,4^. Although eco-hydraulics has achieved successful practices in salmonid fish conservation^5^, dams and water extraction that exacerbate river fragmentation and habitat degradation still account for 46% of fish biodiversity loss^1^. Given the severity of this loss, it is imperative to reconsider what current water resource management practices may be overlooking.

Current research in eco-hydraulics related to fish conservation primarily focuses on river connectivity (e.g., fish passage design^6,7^) and habitat management (e.g., environmental flow theory^8,9^). The underlying concept behind these two approaches is to restore habitats to a semi-natural state (i.e., to improve or maintain river connectivity and habitat availability), thereby supporting sufficiently large populations with adequate genetic diversity to withstand future uncertainties, such as those posed by climate change^10–13^. This concept assumes the ecosystem is static. When the stressors are local — allowing immigrants to offset mortality without altering local genetic diversity, or when biodiversity loss is primarily driven by random genetic drift in small and isolated populations (e.g., due to river fragmentation), such that a large population size and increased population connectivity can buffer against stochastic loss, these two approaches can indeed be effective^13,14^. However, an increasing number of studies show that rapid evolution can happen within one or a few generations^15,16^. For example, reduced river discharge acts as an upper-bound selective pressure, limiting the entry of large-bodied salmon into smaller streams and driving evolutionary shifts toward smaller body sizes^17,18^. Body size is a key parameter influencing swimming ability; thus, it plays a central role in fishway design^5,19^. Fishways function as a lower-bound selective filter by permitting passage only for individuals that exceed a minimum size or swimming threshold. When considering the evolutionary responses in salmon alongside these dual selective pressures on body size — reduced discharge and fishway structure — we can anticipate a long-term decline of average fish body size, potentially rendering future individuals incapable of passing through existing fishways. This joint selective stress on salmon body size could trigger potential ecological feedbacks that deteriorate the ecosystem and its services (e.g., sediment transportation, nutrient transport, fisheries value and meals for rural people)^20,21^.

More generally, the understanding of freshwater fish evolution in response to human perturbation remains limited. Most studies focus on the impacts of human perturbation on genetic diversity from the perspective of river connectivity^3,22–24^, particularly focusing on large barriers^2^, whereas the investigation on the adaptation of freshwater fish has been mainly confined to natural systems, e.g., lake-stream^25^, riffle-pool^26^, waterfalls^27^, and flooding^28^. There are emerging studies on how river impoundment affects fish evolution, but it remains unclear if these changes are plastic or genetic^29^. Besides, biological studies often use categorical variables to describe environmental variation (i.e., spatial^25^ and temporal contrasts^29,30^), overlooking detailed quantitative estimation of the local habitats (e.g., channel cross-section morphology^4^, temporal variance of the discharge^8,31^, or covariance between channel width and discharge^32^). This makes it difficult to connect with the data used by hydraulic engineering. Lastly, extinction in freshwater species is often caused by a combination of factors, such as habitat alterations induced by water management and pollution^1^. While fish have been found to evolve in response to water pollution^33^, physico-chemical conditions and flow regimes are typically studied separately.

The lack of integration between eco-hydraulics and evolutionary biology hinders our understanding of how fish adapt to human perturbations and restricts the applications of evolutionary biology to water resources management. Based on the current gaps we highlighted, this paper aims to transition eco-hydraulics into eco-evo-hydraulics, i.e. a broader framework that also considers rapid evolutionary responses^34^. Here, we focus on two fundamental questions: (1) how does genetic change associate with river habitat in different hydraulic environments, and (2) what functional genes are associated with human perturbation in the river?

We conducted the study in the Demer Basin in Belgium (Figure 1a), which has well-documented physico-chemical monitoring data as well as flow gauging stations, enabling the development of a hydrological model to predict the spatial and temporal variation in discharge^35^. The Demer River is an 85 km rainfed river with a catchment area of 1920 km² and a lowland flow regime. Three-spined stickleback (*Gasterosteus aculeatus* Linnaeus, 1758) (Gasterosteidae, Teleostei) was selected as a model species to study the evolutionary responses of fish to human-regulated flow regimes. Stickleback is a widely used model species in evolutionary biology^36^. Because of their small body size, stickleback are particularly vulnerable to river flow^4^, offering several advantages for examining evolutionary responses^4,3,26,37^. The species also exhibits strong tolerance to pollutants^38,33^, which is important in human-regulated channels that are often contaminated by local industries and agriculture. Freshwater stickleback have limited economic value and are not directly affected by fishing^4^. In 2017, a total of 14 populations were collected from upstream, middle stream and downstream locations of the Demer Basin (Figure 1a; Table S1). Three populations were excluded from the final analysis because of low sample size, high genetic distance to other populations, or lack of hydraulic data (Table S1, Table S2, and Figure S1). Of the 11 remaining populations (genetic diversity summary statistics in Table S3), six populations were located within 200 m downstream of hydraulic barriers (HB populations), and five populations were located in semi-natural streams (SN populations). Semi-natural habitats were channelized over a certain distance with either hard or soft banks, but lacked a hydraulic barrier within 200 m of the sampling sites^4^ (Table S4).

**Figure 1.**
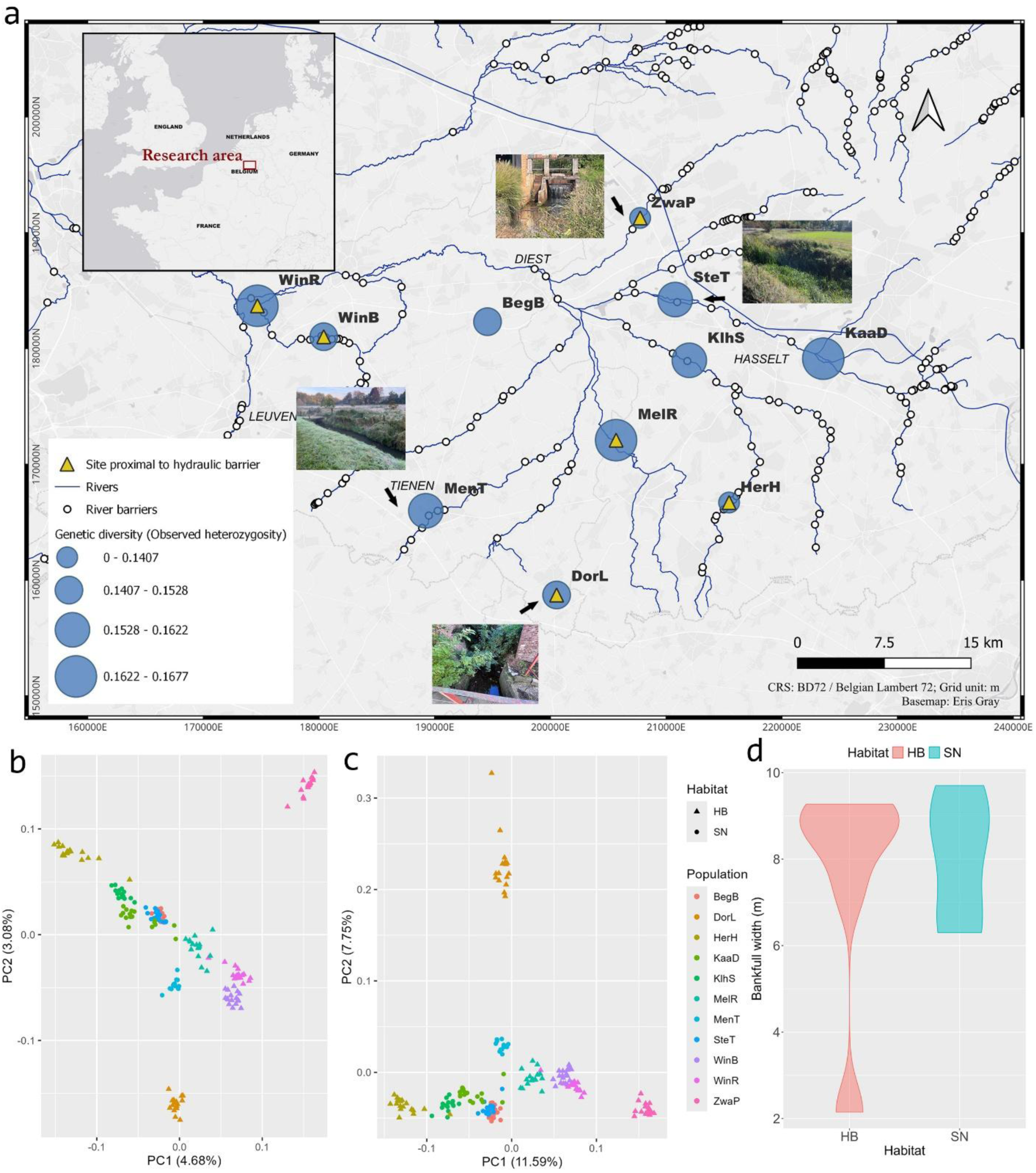
(a) Rivers, hydraulic barriers and genetic diversity (measured as observed heterozygosity, 𝐻_𝑜_) of three-spined stickleback populations sampled at eleven locations. Sampling sites within 200 meter of a hydraulic barriers (HB) are marked with a yellow triangle. Two representative photographs for each habitat type (SN and HB) are shown. (b) The genetic structure of the sampled populations based on principal component analysis (PCA) was calculated with neutral single nucleotide polymorphisms (SNPs) loci and (c) top 5% outlier SNPs based on 𝐹_𝑆𝑇_. (d) Violin plot of the bankfull width of SN and HB sampling sites, showing greater variance in the channelized river. This suggests that river barriers occur across both large and small rivers, with a stronger presence in larger systems.

We combined hydraulic field data with a geological survey, remote sensing, governmental database, basin-wide hydrological modelling^35^, and counts of parasite abundance to compile the following data: (1) river morphology, including sinuosity (-), bed topographical variance (𝑚^2^), relative roughness (*m*), bankfull channel width (*m*) (Figure 1d), and bankfull channel depth (*m*). Here, bankfull refers to data measured at the elevation of the bankfull level, where water starts to overflow into the floodplain. (2) Spatial location, including number of barriers downstream, downstream river distance (*km*), and upstream river distance (*km*). (3) Physico-chemical variables, including average conductivity (𝜇𝑆/𝑐𝑚), average chemical oxygen demand (COD) (*mg/L*), average pH (-), average dissolved oxygen concentration (*mg/L*), and average temperature (℃). (4) River flow regime, including annual averaged daily discharge (𝑚^3^/𝑠), annual averaged unit discharge (𝑚^2^/𝑠) and annual variance of daily discharge (𝑚^6^/𝑠^2^). (5) Parasite abundance, including the abundance of *Gyrodactylus sp.*, *Trichodina sp.* and *Glugea anomala* (descriptive statistics of environmental data in Table S6 and S7). Genotyping-by-sequencing (GBS) was used to identify 69,672 single nucleotide polymorphisms (SNPs) for the calculation of population genetic diversity (Figure 1a) and population genetic structure (Figure 1b-c).

## Intra-riverine allelic frequency changes associated with hydraulic regimes

We first performed redundancy analysis (RDA) on individual genotypes to calculate and compare the explanatory power of each environmental variable separately (Table 1), using adjusted 𝑅^2^ and relative explanatory power index (*REPI*), as detailed in the Materials and Methods. Among all variables, the annual variance of daily discharge showed the strongest association with genetic variance (adjusted 𝑅^2^ = 0.024 and *REPI* = 13.67 %, Table 1). This explanatory power exceeded that of two commonly discussed factors in conservation genomics^3^: the number of barriers downstream (adjusted 𝑅^2^= 0.023 and *REPI* = 12.53%, Table 1) and downstream river distance (adjusted 𝑅^2^ = 0.019 and *REPI* = 10.66%, Table 1). This comparison emphasizes the importance of measuring the local environment and its impact on fish evolution.

**Table 1.** Adjusted 𝑅^2^ and relative explanatory power index (*REPI*) of the correlation of the 19 environmental variables with genetic variance.

| Environmental variable | Adjusted $R^2$ | *REPI* |
| --- | --- | --- |
| Spatial location (dummy variable) | 0.1810 | 100% |
| Annual variance of daily discharge | 0.0247 | 13.67% |
| Annual averaged daily discharge | 0.0243 | 13.44% |
| Annual averaged unit discharge | 0.0237 | 13.12% |
| Number of barriers downstream | 0.0226 | 12.53% |
| Downstream river distance | 0.0193 | 10.66% |
| Average *pH* | 0.0183 | 10.12% |
| Average temperature | 0.0178 | 9.85% |
| Bankfull channel width | 0.0172 | 9.55% |
| Average conductivity | 0.0169 | 9.36% |
| Bankfull channel depth | 0.0146 | 8.11% |
| Bed topographical variance | 0.0145 | 8.02% |
| Relative roughness | 0.0142 | 7.86% |
| Average dissolved oxygen concentration | 0.0137 | 7.58% |
| Upstream river distance | 0.0127 | 7.02% |
| Average *COD* | 0.0124 | 6.85% |
| *Gyrodactylus sp.* abundance | 0.0123 | 6.82% |
| Sinuosity | 0.0099 | 5.51% |
| *Glugea anomala* abundance | 0.0098 | 5.42% |
| *Trichodina sp.* abundance | 0.0093 | 5.15% |

Then, a multi-variable RDA analysis was performed on the individual genotypes to acquire the cumulative explanatory power of all environmental variables simultaneously. To control for collinearity among environmental variables, we applied correlation analysis and step-wise RDA to investigate the variance inflation factors (VIF < 5) (see Materials and Methods). The final model included, ranked from highest to lowest contribution, unit discharge, number of downstream barriers, average pH, longitudinal bed morphological variance, parasite abundance (*Glugea anomala*), and average dissolved oxygen concentration. Collectively, this RDA model constrained 14.7% of the total genetic variance (adjusted 𝑅^2^= 11.1%, *REPI* = 61.3%, Table S10). The model first incorporates variables related to the local hydraulic regime and river fragmentation, followed by variables describing the physico-chemical environment (Table S10, Figure 2a,b). Its explanatory power is significantly higher than models considering only barriers or river distance (Table 1). These findings suggest that local environmental changes could induce fish evolutionary responses, with the hydraulics-related variable, unit discharge, as the most important environmental feature.

**Figure 2.**
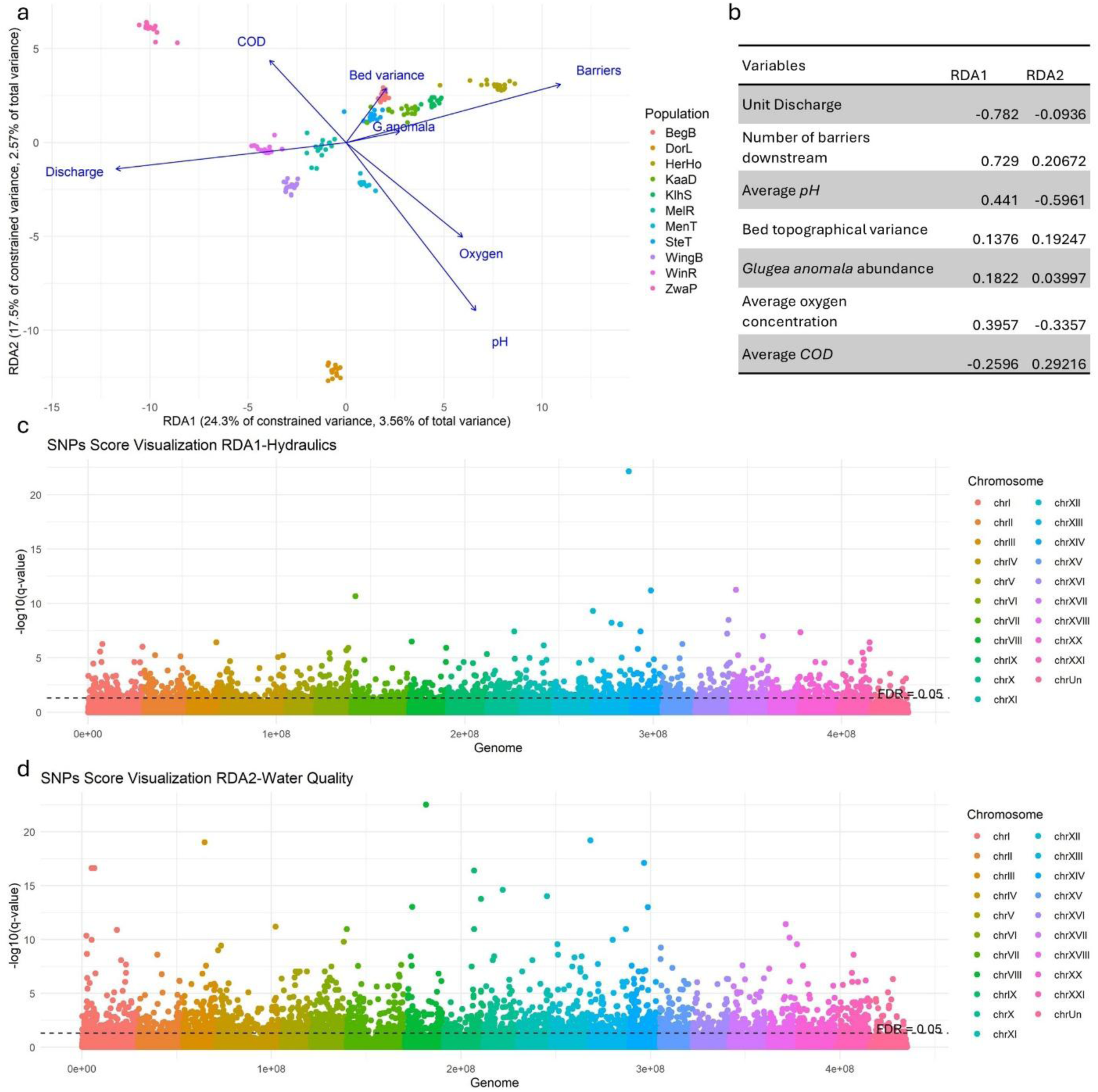
Gene-environmental gradient analysis with redundancy analysis (RDA) and latent factor mixed models (LFMM). (a) Ordination of genetic variance by RDA1 and RDA2 with individuals marked by population. (b) Variables contribution to RDA1 and RDA2. (c) -log10-transformed *q* value for RDA1 of each SNP, arranged by genomic position. (d) -log10-transformed *q* value for RDA2 of each SNP, arranged by genomic position.

## Genes with functions in the nervous and sensory systems correlate with axes of environmental variation

Genotype-environment associations (GxE) were analysed using latent factor mixed models (LFMM)^39^ identifying SNPs that are significantly associated with the two RDA axes acquired above, respectively. In total, 4,137 unique SNPs (5.9% of 69,672 SNPs) were detected across the two RDA axes. RDA1 (1,683 outliers, 2.4% of 69,672 SNPs) was mainly influenced by unit discharge and number of barriers, and RDA2 (2,675 outliers, 3.8% of 69,672 SNPs) was mainly driven by average pH and average dissolved oxygen concentration (Figure 2).

We next assessed the function of the top 10 outliers, as they are more likely to represent true signals, from both RDA axes, based on known gene functions and expression patterns in zebrafish through the Zebrafish (*Danio rerio*) Information Network (ZFIN) (Table S12-S15), where zebrafish serves as a model species to study genomic functions^40^. Based on the underlying functions of genes that are located within 50 kb (Table S12), top outliers for RDA1 are mainly associated with phenotypic responses to environmental stress, particularly involving the nervous, sensory, visual and musculature systems. For instance, *dmrt3a* gene is known for affecting zebrafish swimming ability^41^. In contrast, the top 10 outliers for RDA2 included genes related to molecular functions and regulatory processes. Unlike the outliers for RDA1, only one locus of the top outliers for RDA2 is associated with phenotypes related to the nervous, sensory, and visual systems. While we cannot rule out false positives that merely reflect the action of genetic drift, these functional outliers strengthen the assumption that natural selection acts on specific loci that are related to fish phenotypes that mediate responses to environmental stressors, including hydraulic and hydrological factors.

Enrichment analysis of all the outliers from each RDA provided further insights into the underlying biological processes. RDA1 outliers were primarily involved in biological processes related to signalling and axon guidance, confirming our earlier results that the most significant genes mainly influence neurodevelopment. In contrast, the neurogenesis-related terms of RDA2 fall under cellular component categories (Figure S6-S7). This difference in gene functions suggests that fish use two different mechanisms to regulate their responses to hydraulic/hydrological and physico-chemical stressors, respectively.

## Evolutionary differences between populations with and without a hydraulic barrier

We observed no difference in genetic diversity between HB and SN populations, as indicated by the *U* test (allelic richness, *p-value* = 0.1775; observed heterozygosity, *p-value* = 0.4286), nor in environmental variables of these habitats (Figure 1; Table S7). Gene-habitat interaction analysis using a *Chi-square* test (Figure 3a), least absolute shrinkage and selection operator (LASSO) regression (Figure 3b), and discriminant analysis of principal components (Figure 3c) (Materials and Methods)^42^ identified 372, 72, and 20 divergent SNPs, respectively, between the two habitat types. Among these, a total of 12 outliers were consistently significant across all the tests (Table S16-S17). The 12 outliers were mainly associated with the nervous and sensory systems, aligning with our previous results from the GxE analysis based on LFMM. This finding also aligns with earlier research indicating that sensory systems might influence stickleback swimming behavior^43^.

**Figure 3.**
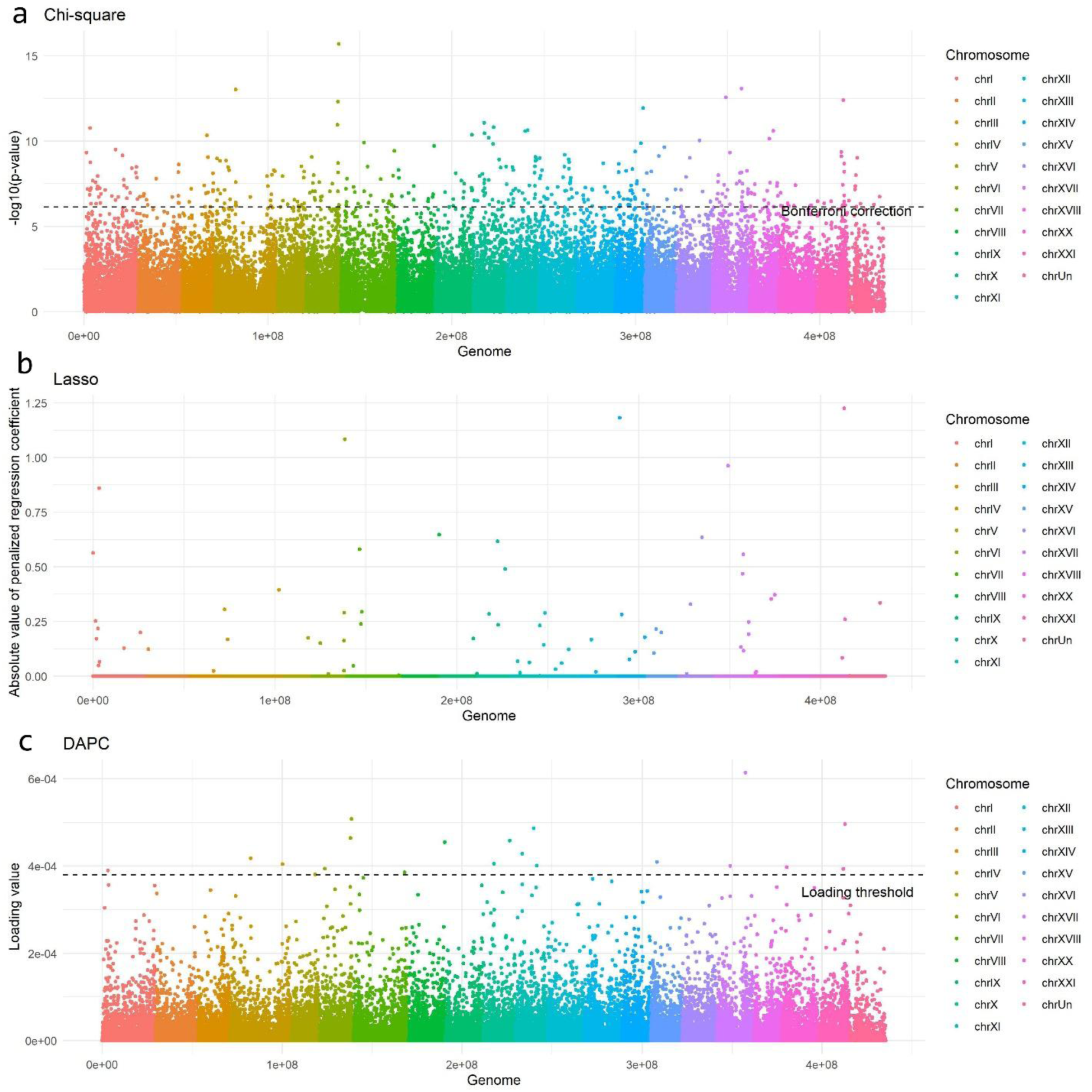
Analysis of populations with and without a hydraulic barrier (a). Chi-square test results. (b). LASSO loading plot. (c). DAPC results.

## Discussion

### Incorporating fish evolutionary responses into water resources management

Why, even within the same species, do some fish adapt more effectively to human disturbances while others do not? Could genetic differences be responsible? Since the average freshwater wildlife population size has dropped by 85% over the last 50 years (1970-2020)^44^, it is crucial to find answers to these questions. For example, if the genetic variation is not observable and instead driven by epigenetic mechanisms^45^, we might still treat the river as a relatively static system in water resources management. However, if genetic differentiation underlies this variation, it becomes necessary to consider the evolutionary dynamics of fish populations into river management (e.g., how local adaptation influences basin-wide adaptation). Identifying the functional genes involved in adaptation could inform alternative conservation strategies—such as improved fish ladder designs, more accurate risk assessments, and targeted habitat protection^34^.

In this study, we analyzed whether or not natural populations of a small riverine fish, three-spined stickleback, show genetic evolutionary responses to intra-riverine human regulation. Our multidisciplinary approach reveals that 5.9% of all the detected loci across the genome show significant associations with basin-wide human-induced variation in river habitat. In addition, we observed an evolutionary response in stickleback fish in the presence of hydraulic barriers. The findings suggest that the nervous and sensory systems are targeted by putative selection in the human-regulated system. Collectively, our result advocates the suggestion of incorporating fish evolutionary responses in eco-hydraulics as a new strategy to address future challenges connected to human impact.

From the application perspective in water resources management, genetic outliers identified by the LFMM model could be used to study evolutionary potential through genomic offset — the distance between the current and required genomic composition in a set of putatively adaptive loci under a future/changed environment ^46^. For instance, the ability of the focal populations to adapt to changes in river hydraulics can be estimated by the genetic diversity of the functional alleles at the loci linked to RDA1. Alternatively, the genetic diversity of all the functional alleles at loci linked to both RDA1 (river hydraulic environment) and RDA2 (river physico-chemical environment) can be used as an indicator for the adaptive potential to river regulation. Combined with basin-wide monitoring data, this RDA model helps to identify the principal environmental drivers of population connectivity, and to improve habitat management. However, because the explanatory power of the RDA model is sensitive to sample size, type and number of genetic markers, and environmental data resolution, we do not recommend cross-comparing the RDA model in different systems unless such sources of bias between studies are properly accounted for.

The set of environmental variables selected in the multi-variable RDA model jointly reflects the range of environmental conditions to which stickleback populations are currently adapted. Genetic variance is found to be associated with unit discharge, which aligns with the natural flow regime^8^. Ignoring spatial-temporal variation in flow regimes could limit our understanding of important environmental determinants of genetic diversity, potentially underestimating the importance of certain hydraulic factors^31^. For example, discharge is an important habitat feature that characterizes if fish experience high or low variation in flow regimes, which potentially affects feeding, shelter, reproduction, exposure to parasites, and other important ecological conditions^8^.

Our study demonstrates that collaboration between water resources management and evolutionary biology can create new opportunities for nature management. The interdisciplinary work of eco-evo-hydraulics can help forecast evolutionary responses of riverine fish to future environmental change^47^. These two fields are frequently dating but not married. Interdisciplinary efforts integrating eco-hydraulics and biology vary across spatial scales. At the local scale, where barriers are constructed, communication and collaboration between eco-hydraulics and biologists are well-established, particularly in fish passage design^24,48,49^. At the basin-wide scale, biological information is commonly collected using genetic tools for monitoring or environmental impact assessment. For example, environmental DNA has been used to monitor species diversity and distribution patterns^50,51^, and genetic markers have been used to assess population genetic structure, which is indicative of the impact of hydraulic infrastructure on river connectivity^3,4,49^. However, integration of hydraulics, hydrology and biology, specifically population genomics, remains limited at the basin-wide scale. This is partly due to uncertainties regarding intra-riverine (basin-wide) adaptive divergence, with currently-known evolutionary changes typically limited to diadromous fish^18^, and primarily focus on conversion from lotic to lentic environments due to reservoir impoundment ^29,52^.

### Evolution to be smarter or stronger?

Our initial assumption was that genetic outliers associated with hydraulic perturbation would primarily involve pathways related to fish anatomical development, such as their fin morphology, which can influence fish swimming ability. However, while some of the top outliers from RDA1 are linked to fish anatomical structures, a larger proportion are associated with the nervous, sensory, and visual systems. These functions may also influence swimming ability but are not easily detectable through conventional phenotypic measurements. This suggests that future studies should also consider fish behaviour or their brain functions^53^, such as how fish navigate in turbulent water. Previous studies have shown that juvenile stickleback exposed to complex learning tasks exhibit temporarily reduced growth, higher mortality and impaired ability to cope with new cognitive challenges later in life^54^. Combined with our findings, this suggests river regulation should also consider reducing perturbations at specific fish life history stages. Understanding how functional genes enhance fish behavior to adapt to different hydraulic perturbations could inform new research in fish passage design.

Our study used genotype-by-sequencing methods rather than whole genome sequencing, so the outliers we identified only represent a fraction of the genome. Consequently, while our dataset is reasonable to test if fish show evolutionary responses, it lacks the resolution to identify large-effect genes that impact fish swimming behavior. Thus, we cannot exclude the important functions of morphologically related genes in fish adaptation. Besides, the LFMM model tends to bias toward genes that exhibit heterozygosity in only a single population. For example, the strongest large-effect locus identified by our LFMM model on the RDA1 axis lies near the *dmrt3a* gene^41^ (Table S12), which is known to affect fish swimming. This large-effect locus could be a true signal. However, it could also be driven by the unique locus polymorphism in the ZwaP sampling site. In the RDA1–RDA2 ordination plot (Figure 2a), ZwaP appears at the extreme end of the RDA1 axis, exhibiting the lowest score along this gradient. As shown in Figure 2a, the environmental vector for discharge is strongly aligned with the negative direction of RDA1, and ZwaP shows the highest loading in that direction. This suggests that ZwaP is particularly affected by discharge-related variation. The strong signal at this single population could therefore reflect a true local adaptive response, especially considering the functional role of the associated gene. However, because the minor allele at this locus was detected only in ZwaP, it may also represent a localized mutation with limited relevance at the broader scale. Nevertheless, if this local mutation indeed confers a functional adaptive advantage in response to anthropogenic disturbance, it raises an important management consideration that enhancing connectivity between this local population and surrounding populations could facilitate the spread of the adaptive allele through gene flow, thereby increasing the overall resilience of the river basin.

In summary, our findings reveal that fish exhibit genetic responses to basin-wide river regulation, underscoring the need for interdisciplinary research between conservation genomics and water resources management. Adaptive capacity can enhance the resilience of water resources management, while techniques in water resources management can in turn provide spatial and temporal data that are challenging to capture through classical methods. This study also outlines that the discovery of fish adaptation to human perturbations could be linked to their neurological and sensory development. We suggest future studies on fish behaviour to explore these possibilities. Finally, we present the potential implications of our results to water resources management and hope to encourage further studies in this emerging field^34^.

## Materials and Methods

### Fish sampling and bioinformatics analysis

Fish were sampled between September and November of 2017 along a 100-meter river section at 14 different locations within the Demer basin in Flanders, Belgium (Figure 1a). A standardized electrofishing method authorized by the Agency of Nature and Forest (ANB) of the Flemish Community was used to collect the fish. Up to 30 three-spined stickleback were collected at each location and euthanized using MS222, following the guidelines of the KU Leuven Animal Ethics Committee. Euthanized individuals were stored at −20°C before fin clips were taken and preserved in 70% ethanol. Any surplus individuals and all other fish were released back into their natural habitat at the sampling location^3^.

### Molecular methods and DNA sequencing

A total of 280 individuals, collected from 14 sampling locations, were genotyped with single nucleotide polymorphisms (SNP) using a Genotyping-by-Sequencing (GBS) protocol as described in Elshire et al. (2011)^55^ and Christiansen et al. (2021)^56^. DNA extractions followed a salt precipitation protocol adapted from Cruz et al. (2017)^57^. DNA quantity was measured using Quant-iT PicoGreen dsDNA kit (Thermo Fisher Scientific Inc.) according to the manufacturer’s instructions. Subsequently, the quality of the extracted DNA was evaluated using agarose gel electrophoresis after diluting the samples to a concentration of 10 ng/μl. A total of three pools, each including 4 replicates and 92 individual GBS libraries, were prepared (92 individuals × 3 + 4 replicates individuals = 280). The extracted DNA was first digested using *ApeKI* at a temperature of 75°C for 2 h. The selection of this restriction enzyme was based on *in silico* optimization, using SimRAD v0.96 ^56,58^ in R v4.0.2 as described by Christiansen et al. (2021)^56^. A barcoded adaptor (4–8 bp) along with a common adaptor were ligated to the digested DNA at 22°C for 60 min, followed by an additional 30 min at 65°C. Subsequently, all samples were purified using CleanPCR (CleanNA) beads, both before and after PCR amplification. The final concentration of each sample was determined using the Quant-iT PicoGreen dsDNA kit (Thermo Fisher Scientific Inc.). Based on individual concentration, samples from each pool were combined to ensure the addition of 10 ng amplified DNA from each individual sample. Each pooled library was subjected to size selection for fragments ranging from 240 to 340 bp (excluding adapters) utilizing a BluePippin (Sage Science). The libraries were 100 bp paired-end sequenced on an Illumina HiSeq2500 platform at Macrogen, Inc.

### Bioinformatics analysis

First, the quality of the raw sequencing reads was assessed using FastQC software v0.11.9^59^. The raw reads were then demultiplexed using the process_radtags module from Stacks v2.54, removing reads with an uncalled base and low-quality scores (Phred < 10)^60–62^. Sequences containing one barcode and/or RAD-Tag mismatch were rescued.

The demultiplexed reads were mapped against the reference genome^63^, using BWA v0.7.17^64^. We remapped the sequence to the stickleback v5 assembly reference genome^63^, which is used by the National Center for Biotechnology Information (NCBI). SNPs were called using GATK v4.2.0.0^65^. Non-variants and non-biallelic SNPs were excluded. Subsequently, SNPs were filtered using VCFtools v0.1.16^66^ and PLINK v2.00a3.1^67^ with the following criteria: mean depth > 10, minor allele count > 3, (genotype) quality > 20, missingness < 20, and heterozygosity > 0.5. Further, linkage disequilibrium was evaluated, and only one SNP was retained if linkage disequilibrium (LD) was detected. Individuals with a depth below 10 and missingness exceeding 20% were excluded. We did not remove SNPs that do not fit with the Hardy–Weinberg equilibrium filter because we also explored the loci that are under selection. In addition, studies show that conducting the Hardy–Weinberg equilibrium filter can impact the inference of population structure^68^. The VCF file was converted to genind file by vcfR v1.15.0 package in R version 4.2.2.

### Genomic data filtering

In one out of 14 locations (FonT, Table S1), the sample size was lower than 10 after bioinformatic filtering, and it was excluded from the data analysis. River morphological and base flow data were collected in the autumn of 2022. We were unable to access one sampling site due to construction (VelG, Table S1), and this location was also excluded from the analysis. After filtering out SNPs on the sex chromosome^69^, a total of 12 populations including 192 individuals and 69,672 SNPs were used to analyze population structure. We included 1,197 SNPs that are on chrUn in the following analysis. Pairwise 𝐹_𝑆𝑇_(Nei 1987^70^) was calculated with the hierfstat (v0.5-11) package in R (Table S2). Averaged pairwise 𝐹_𝑆𝑇_ was calculated, and population DemT (Table S2) was found as an outlier (z-score = 2.46). This high genetic difference of population DemT could be shaped by drift or historical connection between Maas and Scheldt watershed^33^. We, thus, removed DemT population in data analysis. Population structure was calculated with the ‘adegenet’ v2.1.10 package. Using PCA plot, we identified one individual in population DorL that overlapped with BegB and SteT (Figure S1). This might be due to mislabeling. Therefore, we removed this individual for further analysis. Finally, in total 11 populations were used in the analysis (Table S1). Observed heterozygosity, expected heterozygosity, inbreeding coefficient and allelic richness were calculated using the hierfstat (v0.5-11) package in R (Table S3).

### Sampling site description

Focusing on our research question regarding the response of fish to different hydraulic environments, our field survey aimed to provide information for classifying fish habitats based on the intensity of hydraulic impacts. Field observations in the year 2022 were complemented with information from the government website (waterinfo.be) and the GIS distribution of hydraulic barriers (year 2004; http://www.vismigratie.be)^4^.

The sampling sites were classified as hydraulic regulated channels (HB) where we found a hydraulic barrier (low-head dam) within 200 m (Table S4), and semi-natural channels (SN), where channels are channelized or bank reinforced but there is no hydraulic barrier nearby. In Belgium, rarely any river channel has not been exposed to manipulation. Among the HB sites, WinR does not have a typical low-head dam such as a weir or a watermill, but we observed a channelized region with a culvert and water gauge which could disturb the local habitat. This culvert connects Leibeek and Winge streams (Table S4), and the WinR sampling site is under a high probability of flooding (waterinfo.be). Frequent flooding makes a culvert behave as a hydraulic barrier, leading to a high probability of disturbing the local hydraulic habitat. There are six HB channels and five SN channels.

### Environmental Data Collection

#### Physio-chemical data

Water physio-chemical averaged data, including average conductivity (𝜇𝑆/𝑐𝑚), average chemical oxygen demand (COD) (*mg/L*), average pH (-), average dissolved oxygen (*mg/L*), and average temperature (℃), were downloaded from the Flemish Environment Agency (VMM) website for the year 2017.

#### Parasite abundance

We monitored the abundance of three ectoparasites (*Gyrodactylus spp.*, *Trichodina sp.* and *Glugea anomala*) because parasites represent major agents of selection in stickleback^71^ and may also impact fish swimming behavior^72^. Nine populations were screened by Deflem et al. (2022)^73^; two additional populations were measured in 2022 following the same protocol. Fish at the other two populations were cut by one side of the pectoral fin and the caudal fin for sequencing, while the remaining body was used for parasite abundance measurements. First, we rehydrated the fish in distilled water in 5 or 10 ml cryo-tubes with 1 or 2 ml of distilled water. After a vigorous shake of 10 s, the liquid was poured into a Petri dish for parasite screening. Because one pectoral fin and the caudal fin were cut, we used the parasite abundance of *Gyrodactylus* at different regions of fish^74^ to correct the abundance of data. One pectoral fin and the caudal fin account for 10% and 7% of *Gyrodactylus* abundance, respectively^74^. So, we multiply the parasite abundance data by 1.17.

#### Channel morphology

The total number of barriers to the river mouth, downstream distance and upstream distance was acquired from VMM measured or counted from the mouth of the Demer as the most downstream location (50°58’07.1“N 4°41’33.8”E), and the bankfull width and bankfull depth were measured in October 2022 on site. Sinuosity was measured from QGIS based on remote sensing data from OpenStreetMap along rivers measured by the same person. Channel sections and longitudinal bed elevation profiles were measured in August 2023. We initially used Real Time Kinematic Global Positioning System (RTK-GPS) with the mount point provided by the Agency Digital Flanders (FLEPOSVRS32GREC). However, some of our sampling sites had high tree canopies, and some locations had base stations, which disturbed our RTK-GPS signal. We could only perform RTK-GPS measurements at seven sites, and the remaining sites were measured with classical methods that measure water depth at equal intersections (an arm length of the same person). Because all sites were measured by the same person at the same length of intersections, we calculated the topological variance at the same weight of each sampling data. At DorL and SteT, we could not enter the river because of construction and dense vegetation. Therefore, we used the bed roughness at a similar habitat for this location (Table S5). Finally, relative roughness (topological variance/bankfull depth) was computed by using longitudinal topological elevation variance divided by bankfull depth. Thus, the bed roughness, represented by the bed elevation variance, is a semi-quantitative way to describe habitats, including vegetation and bed substrate.

#### Flow regime modelling and calibration

For each sample location the unit discharge (discharge/bankfull width), the annual daily average discharge and variance were used. These variables are calculated on the daily river discharges which are obtained by applying a non-parametric data-driven approach based on spatial interpolation. This method has been applied to the Flanders region of Belgium, and has proven to be as good as two state-of-the art hydrological models^35^. The applied method takes into account the anthropogenic influences as reflected in the observations, which can be of high relevance in areas with a lot of hydraulic disturbances such as the Demer basin.

In order to calibrate the non-parametric data-driven approach to our sampling sites, the surface velocity at each site was measured with a floater between 10^th^-13^th^ Oct 2022. There was no intensive rainfall one week before our sampling date and during our sampling period. Based on the measured velocity, the river discharge is calculated by applying the velocity-area method with a coefficient of 0.85 to convert surface velocity to depth-averaged velocity^75^. There were considerable challenges in measuring surface velocity in these channels such as vegetation coverage. However, given that the applied non-parametric data-driven approach has proven to be as good as state-of-the art hydrological models in Flanders and that the measurements are used to further optimize the results, the impact of measurement uncertainties on the end results are assumed to be minimal.

### Statistic analysis

#### Genetic diversity and population structure

Observed heterozygosity (*H_o_*), expected heterozygosity (*H_e_*), inbreeding coefficient (*F_IS_*), and allelic richness (AR) were calculated with hierfstat v0.5-11 in package in R (Table S3). Population structure was calculated using principal component analysis (PCA) with ggfortify v0.4.17 package in R (Figure 1b-c). Population structure was also calculated by applying Non-metric MultiDimensional Scaling (NMDS) to the pairwise 𝐹_𝑆𝑇_ data with the vegan v4.2.3 package in R (Figure S2).

#### Environmental data descriptive analysis

Environmental data were checked for outliers through a z-score [-3, 3], followed by descriptive statistics through R package psych v2.4.6.26, and Shapiro Wilk’s *W* test to check dataset normality through R package stats v4.2.2. The results are shown in Table S6.

Descriptive statistics were applied within each of the two categories of habitats, i.e. sites with a hydraulic barrier (HB) and semi-natural (SN) sites. Whitney *U* test (stats v4.2.2) was applied to each variable to test if there was a significant difference. The results are listed in Table S7.

#### Redundancy analysis on the explanation power of environmental variables

Redundancy analysis (RDA) combines ordination and regression on two or more data frames, and it has been widely used in genotype-environment association (GEA) analysis to study the explanatory power of multiple environmental variables to multiple genotypes^76^. Here, we used R package vegan v2.6-4 that is commonly used in GEA. The output of RDA, as in other regression models, is the (adjusted) coefficient of determination (𝑅^2^). However, it is rarely the case in population genetics that the sample size of each population and the number of SNPs identified remain consistent, whether in a single study or across multiple studies. Thus, the value of the adjusted-𝑅^2^ is difficult to compare among studies. Therefore, we proposed a relative explanatory power index (*REPI*), which normalizes the adjusted R-square of a model with environmental variables to spatial location as a dummy variable.

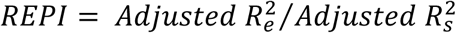

> Where, 𝐴𝑑𝑗𝑢𝑠𝑡𝑒𝑑 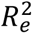 = adjusted coefficient of determination of the RDA model with environmental variables as explanatory variables and genotypes (SNPs allelic frequency here) as the response variable;
>
> 𝐴𝑑𝑗𝑢𝑠𝑡𝑒𝑑 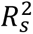 = adjusted coefficient of determination of RDA model with spatial location as explanatory variables (dummy variables) and genotypes (SNPs allelic frequency here) as the response variable.We first compared the explanatory power of hydrological data (temporal average and temporal variance of discharge) from the fish sampling year 2017 (short-term) with the hydrological data collected for the years 2015 to 2017 (long-term). The comparison showed that short-term hydrological data from the 2017 fish sampling year explained more genetic variance than the long-term averaged data (adjusted 𝑅^2^, Table S8). The short-term hydrological data from the sampling year was then used for the following analysis.

To avoid collinearity of environmental variables, we conducted multiple analyses. First, we conducted a correlation analysis for all the environmental variables (Table S9). From the correlation results, we identified a strong correlation in hydrological variables. High correlations in hydrological variables are found at *Annual averaged daily discharge ∼ Annual variance of daily discharge* (*r* = 0.95, *p* < 0.01). In order to decide which variable to keep, we applied RDA analysis on each variance to the genetic variance and compared their explanatory power (Table 1). We kept the corrected temporal discharge variance because of its higher explanatory power. In the empirical approach in hydraulic engineering, (bankfull) channel width is often described by a power function of (bankfull) discharge^32^. Here, we transformed the corrected average discharge to unit discharge by dividing it by bankfull channel width. This yielded an additional variable, which can indicate the vertical velocity profile of the channel. Through this transformation, we reduce the correlation between the magnitude of discharge and variance of discharge: *Annual averaged daily discharge ∼ Annual averaged unit discharge* (*r* = 0.84, *p* < 0.01).

In the channel morphological data, we found another high correlation between *Bed topographical variance (variance of longitudinal bed elevation) ∼ Bankfull channel depth* (*r* = 0.85, *p* < 0.01). Considering that morphological longitudinal variance can describe river habitat, which is independent of bankfull channel depth, we kept both variables and used the variance inflation factor (VIF) at RDA analysis to verify the level of collinearity. We also calculated relative roughness, which is a variable calculated by using morphological longitudinal variance divided by bank full depth to scale the impacts from different sizes of channels. However, because the sampling channels are channelized channels, we found an even stronger correlation in bed morphology *Bed topographical variance ∼ Relative roughness* (*r* = 0.99, *p* < 0.01). Thus, we removed the relative roughness in the subsequent RDA analysis.

The remaining variables were used in a forward stepwise RDA analysis to reduce dimensionality, and the RDA output model was tested with criterion, VIF < 5, to control collinearity (Table S10). We removed the variable with the highest VIF value and conducted the stepwise model with the remaining variables from the previous step until the VIF value of all the selected variables satisfied the VIF criterion. While rerunning the stepwise model does not alter the VIF values of variables in the subsequent model, it helps to clarify the ranking of each variable by making their contributions to the model easier to interpret. Our final RDA output (Table S10-S11, adjusted 𝑅^2^ = 0.111, *REPI* = 61.2%) included, in order of explanatory power: unit discharge, number of barriers to downstream, average pH, morphological longitudinal variance, parasite abundance (*G. anomala*) and average dissolved oxygen concentration. We also tested a partial RDA model with downstream distance as a control variable. However, this partial RDA model did not increase model explanation power (adjusted 𝑅^2^ = 0.102, *REPI* = 56.4%) and downstream distance conflicted with our VIF criterion (Table S10).

#### Gene-Environmental Association Studies with LFMM

Gene-environmental association analysis (GxE) was conducted with the LFMM model^39,77^ using the R package lfmm v1.1. First, the integer for the number of latent factors (*K* = 2) in the LFMM model was jointly identified using a scree plot of the multi-variable RDA output and the PCA of the genetic data (Figure S3). Then LFMM was applied with a ridge penalty^39^. The significant value of the LFMM model was calibrated by the genome inflation factor (GIF). By observing the histogram frequency of the first three RDA axes (Figure S4), the first two RDA axes were identified as selection signals, which were used to identify genetic outliers. The calibrated *p*-values were converted to *q* value with R package qvalue v2.30.0, and we set the false discovery rate to *q*-value = 0.05.

Following the identification of outliers of the two RDA axes, we conducted two types of analysis. First, the top 10 significant SNPs were used to identify genes and functions from NCBI Genome Data Viewer (RefSeq assembly GCF_016920845.1). If an SNP is located within a coding gene (or a non-coding RNA), we use the gene/non-coding RNA to search for the function. Otherwise, we took the nearest two coding genes at each side of the SNP. The gene functions, expressions, and associated phenotypes were collected from the Zebrafish Information Network (https://zfin.org). If a polymorphism only occurred in one population that has a high value in the RDA axes, it can induce a highly significant result. Therefore, we further checked the number of populations that are polymorphic at that SNP (Table S12-S15).

Second, enrichment analysis was used to show the gene functional pathways for all the outliers of each RDA axis and all the outliers for both axes through the g:Profiler online platform^78^. In order to perform the enrichment analysis, we used the rtracklayer v1.58.0 and GenomicRanges v1.50.2 packages to extract the protein id from the stickleback Ensembl annotations (release 95). This analysis only retained the SNPs located within a gene. G:Profiler used the parameters: organism ‘*Gasterosteus aculeatus* (stickleback)’, statical domain ‘only annotated genes’, significant threshold ‘g:SCS threshold’, threshold ‘0.05’ (Figure S5-S7).

#### Categorical gene-environmental analysis on intensively regulated habitat and semi-natural habitat

Populations from the two habitat types are labelled in the PCA plot (Figure 1b-c) and NMDS plot (Figure S2), which show consistent population structure for the two methods. A Whitney *U* test between the two habitat types was applied to the population allelic richness; the result was not significant (*p-value* = 0.1775).

Next, we applied a Chi-square analysis to test if the genotype of each SNP differed between the two habitats (SN and HB). P-values of the Chi-square test were corrected with qvalue v2.30.0 package in R and we set the false discovery rate to 0.05. There were 15,175 SNPs that showed a significant contrast. For comparison, we also used Bonferroni correction on the p-value. Only 372 SNPs showed significance (Figure 3a).

Third, the least absolute shrinkage and selection operator (LASSO) regression model (family = “binomial”, lambda.min.ratio=0.01, alpha = 1), was performed to identify SNPs that can be used to classify two habitats. This analysis was done with R package glmnet v4.1-8, and revealed 72 outliers (Figure 3b).

Fourth, we performed discriminant analysis of principal components (DAPC) in R package adegenet v2.1.10 as another way to find SNPs that differentiate the individuals from the two habitats. Cross-validation retained 40 PCAs for DAPC analysis, and because this is a classification problem on one criterion, we set n.da = 1. Finally, 20 SNPs were identified and showed a good performance in classifying two habitats (Figure 3c and Figure S8). There were 12 SNPs that showed consistent significant divergence across the three analyses. We used the same procedures as the top genes in the GxE analysis in the environmental gradient to search for the function and expression as well as associated phenotypes from the databases (Table S16-S17).

## Acknowledgements

We would like to extend our acknowledgement to Steven Van Belleghem, Tim Maes, Hugo Gante and Jimmy Van Criekingen at KU Leuven for their help with sampling and data analysis. We also thank uOGlobal Power Corporation of Canada Scholarships and the University of Ottawa International Experience Scholarships grant to Cai, and the Natural Sciences and Engineering Research Council (NSERC) Discovery grant to Rennie.

## CRediT authorship contribution statement

**Xiatong Cai and Io Deflem equally** contributed to Conceptualization, Methodology, Validation, Formal analysis, Investigation, Writing - Original Draft, Writing - Review & Editing, and Visualization throughout the whole project. **Joke De Meester and Patrick Willems** contributed to Methodology, Validation, Formal analysis, Investigation, Writing - Original Draft, and Writing - Review & Editing of the hydrological simulation section. **Bart Hellemans** supervised the fish genotyping and contributed to Methodology, Investigation, and Writing - Review & Editing. **Federico Calboli and Andrew Hendry** contributed to Conceptualization, Methodology, Investigation, Writing - Review & Editing. **Colin D. Rennie, Joost A.M. Raeymaekers and Filip A.M. Volckaert** contributed to Conceptualization, Methodology, Investigation, Resources, Writing - Review & Editing, Project administration, Supervision, and Funding acquisition.

