## Supplemental Materials for "Fish Show Genetic Evolutionary Responses to River Regulation"

### Tables

**Table S1** List of sampling sites with code, coordinate, river and location.

| PopID | Latitude | Longitude | Code | River | Location | Note |
|---|---|---|---|---|---|---|
| GA04 | 50.73678 | 5.0847 | DorL | Dormaalbeek | Landen | Intensively regulated |
| GA10 | 50.96292 | 4.72003 | WinR | Winge | Rotselaar | Intensively regulated |
| GA13 | 50.93865 | 4.80135 | WinB | Winge | Blauwmolen | Intensively regulated |
| GA15 | 50.9494 | 5.002917 | BegB | Begijnenbeek | Bekkevoort | Semi-natural |
| GA20 | 51.02876 | 5.192059 | ZwaP | Zwarte beek | Paal | Intensively regulated |
| GA23 | 50.9651 | 5.23407 | SteT | Steenlaak | Thiewinkel | Semi-natural |
| GA24 | 50.85647 | 5.15955 | MelR | Melsterbeek | Runkelen | Intensively regulated |
| GA25 | 50.91782 | 5.25058 | KlhS | Kleine Herk | Stevoort | Semi-natural |
| GA26 | 50.91774 | 5.41525 | KaaD | Kaatsbeek | Diepenbeek | Semi-natural |
| GA29 | 50.85626 | 4.95008 | VelG | Velpe | Glabbeek | Not accessible in 2022 |
| GA30 | 50.80299 | 4.92535 | MenT | Mene | Tienen | Semi-natural |
| GA45 | 50.81631 | 5.5045 | DemT | Demer | Tongeren | Strongly genetically differentiated |
| GA46 | 50.7962 | 5.4194 | FonT | Fonteinbeek | Tongeren | Low sample size |
| GA47 | 50.80706 | 5.29728 | HerH | Herk | Hoepertingen | Intensively regulated |

**Table S2** Pair-wise $F_{ST}$ values between samples. The stronlgy differentiated GA45 sample is highlighted. For sample site abbreviations see Table S1.

| Population | GA04 | GA10 | GA13 | GA15 | GA20 | GA23 | GA24 | GA25 | GA26 | GA30 | GA45 |
|---|---|---|---|---|---|---|---|---|---|---|---|
| GA10 | 0.0185 | | | | | | | | | | |
| GA13 | 0.0183 | 0.0078 | | | | | | | | | |
| GA15 | 0.0208 | 0.016 | 0.0168 | | | | | | | | |
| GA20 | 0.0286 | 0.019 | 0.0202 | 0.0262 | | | | | | | |
| GA23 | 0.0184 | 0.0129 | 0.0139 | 0.0133 | 0.0234 | | | | | | |
| GA24 | 0.017 | 0.0095 | 0.011 | 0.0142 | 0.0204 | 0.0113 | | | | | |
| GA25 | 0.0181 | 0.015 | 0.015 | 0.014 | 0.025 | 0.0104 | 0.0118 | | | | |
| GA26 | 0.0173 | 0.0142 | 0.0147 | 0.0135 | 0.0246 | 0.0058 | 0.0116 | 0.0087 | | | |
| GA30 | 0.017 | 0.0139 | 0.0132 | 0.0171 | 0.0247 | 0.0151 | 0.0133 | 0.0132 | 0.0141 | | |
| GA45 | **0.0321** | **0.035** | **0.0341** | **0.0333** | **0.0427** | **0.0279** | **0.0315** | **0.0268** | **0.0226** | **0.0314** | |
| GA47 | 0.0227 | 0.0226 | 0.0223 | 0.0184 | 0.0315 | 0.0157 | 0.0176 | 0.0069 | 0.0136 | 0.0175 | 0.0299 |

**Table S3** Table with population genetic diversity, including sample size, observed ($H_o$) and expected ($H_e$) heterozygosity, inbreeding coefficient $F_{IS}$ and allelic richness (AR). For sample site abbreviations see Table S1.

| Population | Sample size | *Hobs* | *Hexp* | $F_{IS}$ | *AR* |
|---|---|---|---|---|---|
| GA04 | 16 | 0.147628 | 0.15083 | 0.021009 | 1.27332 |
| GA10 | 17 | 0.167733 | 0.172222 | 0.025189 | 1.314018 |
| GA13 | 17 | 0.151354 | 0.167097 | 0.078825 | 1.303782 |
| GA15 | 16 | 0.152828 | 0.162376 | 0.054351 | 1.294957 |
| GA20 | 16 | 0.129551 | 0.139634 | 0.059158 | 1.252717 |
| GA23 | 17 | 0.162193 | 0.171897 | 0.048493 | 1.313256 |
| GA24 | 16 | 0.165641 | 0.172343 | 0.033515 | 1.313964 |
| GA25 | 14 | 0.158938 | 0.175571 | 0.076943 | 1.319386 |
| GA26 | 17 | 0.163208 | 0.175322 | 0.058319 | 1.319257 |
| GA30 | 13 | 0.160488 | 0.171503 | 0.052677 | 1.311639 |
| GA47 | 18 | 0.140685 | 0.157496 | 0.091845 | 1.285545 |
| Overall | 177 | 0.1546 | 0.1651 | 0.0638 | 1.300167 |

**Table S4** Classification and description of sampling sites, including site ID, site code, presence of an artificial barrier, river distance to the upstream artificial barrier, type of barrier, river distance to downstream barrier and type of barrier.

| Site ID | Site Code | Whether close to an artificial barrier | River distance to the upstream artificial barrier (m) | Type of barrier | Photograph |
|---|---|---|---|---|---|
| ID04 | DorL | Yes | <20 | A field survey found a gate upstream. | 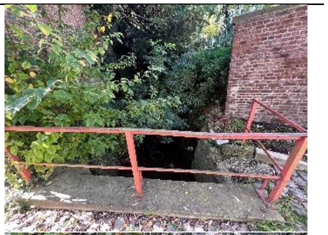 |
| ID10 | WinR | Yes | 124 | A culvert is observed near the sampling site. 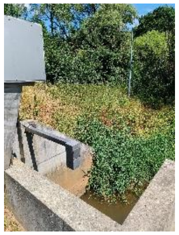 | 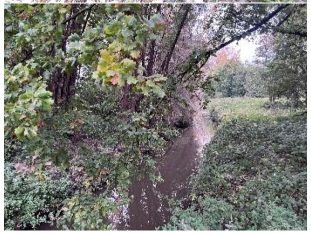 |
| ID13 | WinB | Yes | 152 | A hydraulic structure (water mill) was observed near the road, and only 20 m from the sampling reach. The distance is measured from the mill to the fish sampling point coordinate. | 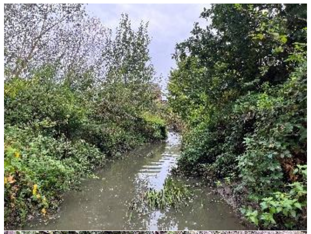 |
| ID15 | BegB | No | N/A | Upstream is a lentic system created by a natural log jam upstream < 20 m | 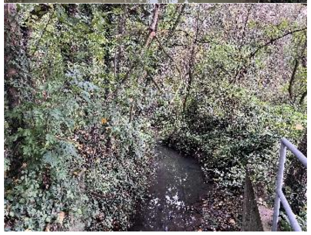 |

| | | | | | |
|---|---|---|---|---|---|
| ID20 | ZwaP | Yes | <50 | Water mill upstream from field survey. | 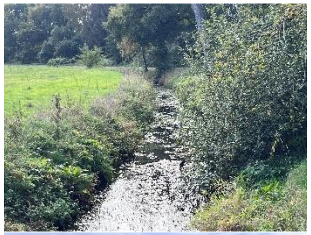 |
| ID23 | SteT | No | N/A | No information from waterinfo.be and GIS layer. We did not find an artifical barrier. | 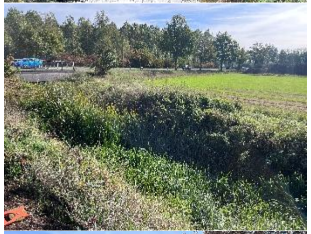 |
| ID24 | MelR | Yes | <50 | No record on GIS and waterinfo.be, but we were informed by local people a watermill is upstream. | 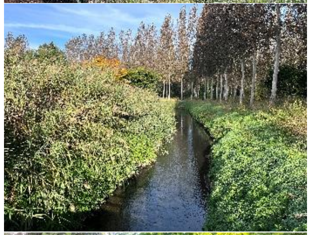 |
| ID25 | KlhS | No | N/A | No information from waterinfo.be and a GIS layer. We did not find an artificial barrier. | 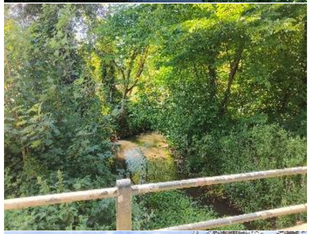 |
| ID26 | KaaD | No | N/A | No information from waterinfo.be and a GIS layer. We did not find an artificial barrier. | 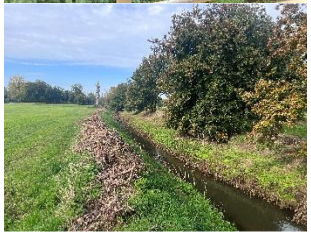 |

| | | | | | |
|---|---|---|---|---|---|
| ID30 | MenT | No | N/A | No information from waterinfo.be and GIS layer. We did not find an artificial barrier. | 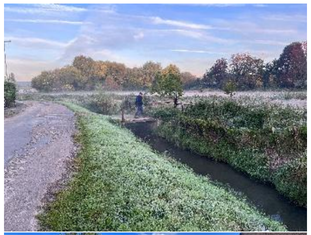 |
| ID47 | HerH | Yes | <20 | We observed fish ladders upstream. | 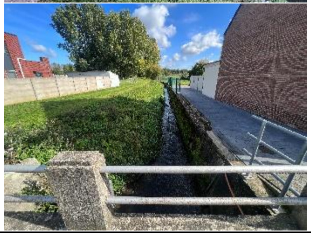 |

**Table S5** Congruence table between the site codes for fish sampling and topological measurements, including the method used.

| Sites ID | Code | Methods |
|---|---|---|
| GA04 | DorL | Used ID 25 data |
| GA10 | WinR | Measured with gauge |
| GA13 | WinB | Measured with RTK GPS |
| GA15 | BegB | Measured with gauge |
| GA20 | ZwaP | Measured with RTK GPS |
| GA23 | SteT | Used ID 26 data |
| GA24 | MelR | Measured with RTK GPS |
| GA25 | KlhS | Measured with RTK GPS |
| GA26 | KaaD | Measured with gauge |
| GA30 | MenT | Measured with RTK GPS |
| GA47 | HerH | Measured with RTK GPS |

**Table S6** Descriptive statistics of the 19 environmental variables documented, including average, standard deviation, median, minimum, maximum, range, skew, kurtosis, standard error and significance of Shapiro's test.

| Variables | mean | standard deviation | standard error | median | min | max | range | skew | kurtosis | Shapiro test (p) |
|---|---|---|---|---|---|---|---|---|---|---|
| **Bankfull channel width** | 7.747 | 2.158 | 0.651 | 8.700 | 2.150 | 9.700 | 7.550 | -1.475 | 1.255 | 0.004 |
| **Bankfull channel depth** | 2.030 | 0.307 | 0.093 | 1.950 | 1.655 | 2.795 | 1.140 | 1.199 | 0.827 | 0.078 |
| **Sinuosity** | 1.171 | 0.099 | 0.030 | 1.139 | 1.065 | 1.410 | 0.345 | 1.132 | 0.359 | 0.048 |
| **Number of barriers downstream** | 13.909 | 7.300 | 2.201 | 14.000 | 1.000 | 30.000 | 29.000 | 0.373 | 0.175 | 0.433 |
| **Downstream river distance** | 46.188 | 21.841 | 6.585 | 48.930 | 2.476 | 68.504 | 66.029 | -0.776 | -0.739 | 0.086 |
| **Upstream river distance** | 15.207 | 7.533 | 2.271 | 13.335 | 6.098 | 29.956 | 23.858 | 0.570 | -1.182 | 0.130 |
| **Average conductivity** | 529.102 | 236.253 | 71.233 | 470.330 | 219.080 | 883.670 | 664.590 | 0.247 | -1.663 | 0.227 |
| **Average COD** | 20.353 | 5.967 | 1.799 | 20.500 | 11.330 | 29.330 | 18.000 | 0.166 | -1.405 | 0.453 |
| **Average pH** | 7.636 | 0.336 | 0.101 | 7.662 | 7.025 | 8.050 | 1.025 | -0.352 | -1.347 | 0.534 |
| **Average oxygen concentration** | 8.764 | 0.758 | 0.228 | 8.910 | 7.630 | 9.850 | 2.220 | 0.021 | -1.674 | 0.484 |
| **Average temperature** | 12.968 | 1.018 | 0.307 | 12.770 | 11.430 | 14.830 | 3.400 | 0.107 | -0.961 | 0.617 |
| **Bed topographical variance** | 0.011 | 0.013 | 0.004 | 0.004 | 0.003 | 0.045 | 0.042 | 1.645 | 1.559 | 0.000 |
| **Annual variance of daily discharge** | 0.015 | 0.020 | 0.006 | 0.007 | 0.001 | 0.068 | 0.067 | 1.646 | 1.771 | 0.001 |
| **Annual averaged daily discharge** | 0.154 | 0.125 | 0.038 | 0.102 | 0.029 | 0.484 | 0.455 | 1.529 | 1.600 | 0.006 |
| **Annual averaged unit discharge** | 0.022 | 0.018 | 0.005 | 0.018 | 0.003 | 0.066 | 0.063 | 1.292 | 0.620 | 0.017 |
| **Gyrodactylus sp. abundance** | 0.114 | 0.109 | 0.033 | 0.100 | 0.000 | 0.334 | 0.334 | 0.588 | -0.990 | 0.248 |
| **Trichodina sp. abundance** | 22.620 | 26.380 | 7.954 | 14.320 | 0.230 | 80.300 | 80.070 | 1.009 | -0.422 | 0.020 |
| **Glugea anomala abundance** | 0.076 | 0.128 | 0.039 | 0.000 | 0.000 | 0.420 | 0.420 | 1.675 | 1.839 | 0.000 |
| **Relative roughness** | 0.005 | 0.005 | 0.001 | 0.002 | 0.001 | 0.016 | 0.015 | 1.276 | 0.281 | 0.001 |

**Table S7** Descriptive statistics of 19 environmental variables for the sampling sites grouped in hydraulic barrier (HB) and semi-natural (SN) channels, including average, standard deviation, median, minimum, maximum, range, skew, kurtosis, standard error and significance of Shapiro's test

| Variables | Classification | mean | standard deviation | standard error | median | min | max | range | skew | kurtosis | Whitney *U* test (p) |
|---|---|---|---|---|---|---|---|---|---|---|---|
| **Bankfull channel width** | HB | 7.520 | 2.720 | 1.111 | 8.650 | 2.150 | 9.270 | 7.120 | -1.181 | -0.412 | 1.000 |
| | SN | 8.020 | 1.489 | 0.666 | 8.700 | 6.300 | 9.700 | 3.400 | -0.147 | -2.131 | |
| **Bankfull channel depth** | HB | 1.987 | 0.215 | 0.088 | 1.965 | 1.655 | 2.270 | 0.615 | -0.156 | -1.489 | 1.000 |
| | SN | 2.081 | 0.414 | 0.185 | 1.950 | 1.750 | 2.795 | 1.045 | 0.885 | -1.135 | |
| **Sinuosity** | HB | 1.199 | 0.120 | 0.049 | 1.176 | 1.065 | 1.410 | 0.345 | 0.618 | -1.142 | 0.429 |
| | SN | 1.137 | 0.061 | 0.027 | 1.113 | 1.091 | 1.240 | 0.149 | 0.854 | -1.205 | |
| **Number of barriers downstream** | HB | 12.667 | 9.893 | 4.039 | 12.000 | 1.000 | 30.000 | 29.000 | 0.574 | -1.072 | 0.247 |
| | SN | 15.400 | 2.408 | 1.077 | 16.000 | 12.000 | 18.000 | 6.000 | -0.289 | -1.871 | |
| **Downstream Distance** | HB | 41.057 | 27.860 | 11.374 | 47.681 | 2.476 | 68.504 | 66.029 | -0.312 | -1.903 | 0.931 |
| | SN | 52.344 | 11.638 | 5.205 | 48.930 | 38.509 | 66.988 | 28.479 | 0.118 | -2.005 | |
| **Upstream Distance** | HB | 16.440 | 6.598 | 2.693 | 17.431 | 6.098 | 22.376 | 16.278 | -0.381 | -1.728 | 0.429 |
| | SN | 13.727 | 9.079 | 4.060 | 9.908 | 9.129 | 29.956 | 20.827 | 1.069 | -0.925 | |
| **Average conductivity** | HB | 521.140 | 281.489 | 114.917 | 463.125 | 219.080 | 883.670 | 664.590 | 0.249 | -1.939 | 1.000 |
| | SN | 538.656 | 200.710 | 89.760 | 470.330 | 340.710 | 755.500 | 414.790 | 0.180 | -2.211 | |
| **Average COD** | HB | 18.702 | 6.434 | 2.627 | 17.710 | 11.330 | 29.330 | 18.000 | 0.445 | -1.423 | 0.429 |
| | SN | 22.334 | 5.317 | 2.378 | 21.000 | 15.500 | 29.000 | 13.500 | 0.024 | -1.898 | |
| **Average pH** | HB | 7.666 | 0.362 | 0.148 | 7.750 | 7.025 | 7.986 | 0.961 | -0.706 | -1.172 | 0.792 |
| | SN | 7.600 | 0.339 | 0.152 | 7.471 | 7.229 | 8.050 | 0.821 | 0.233 | -2.002 | |
| **Average oxygen concentration** | HB | 8.867 | 0.926 | 0.378 | 8.920 | 7.630 | 9.850 | 2.220 | -0.125 | -2.066 | 0.537 |
| | SN | 8.640 | 0.572 | 0.256 | 8.910 | 7.990 | 9.280 | 1.290 | -0.151 | -2.144 | |
| **Average temperature** | HB | 13.098 | 1.169 | 0.477 | 12.810 | 11.490 | 14.830 | 3.340 | 0.173 | -1.528 | 0.662 |
| | SN | 12.812 | 0.909 | 0.406 | 12.770 | 11.430 | 13.700 | 2.270 | -0.408 | -1.657 | |
| **topologicalSpatial Variance** | HB | 0.007 | 0.008 | 0.003 | 0.004 | 0.003 | 0.023 | 0.020 | 1.344 | -0.110 | 0.463 |
| | SN | 0.015 | 0.017 | 0.008 | 0.012 | 0.003 | 0.045 | 0.042 | 0.893 | -1.130 | |

| | | | | | | | | | | | |
|---|---|---|---|---|---|---|---|---|---|---|---|
| **correctedQ TemporalVariance** | HB | 0.021 | 0.025 | 0.010 | 0.014 | 0.002 | 0.068 | 0.065 | 0.966 | -0.756 | 0.247 |
| | SN | 0.007 | 0.009 | 0.004 | 0.003 | 0.001 | 0.023 | 0.022 | 0.917 | -1.126 | |
| **correctedAve Discharge** | HB | 0.206 | 0.146 | 0.060 | 0.170 | 0.091 | 0.484 | 0.393 | 0.983 | -0.692 | 0.082 |
| | SN | 0.092 | 0.057 | 0.026 | 0.083 | 0.029 | 0.183 | 0.154 | 0.508 | -1.449 | |
| **Annual averaged unit discharge** | HB | 0.031 | 0.021 | 0.008 | 0.023 | 0.010 | 0.066 | 0.056 | 0.679 | -1.268 | 0.082 |
| | SN | 0.012 | 0.006 | 0.003 | 0.012 | 0.003 | 0.021 | 0.018 | 0.103 | -1.455 | |
| **Gyrodactylus sp. abundance** | HB | 0.094 | 0.103 | 0.042 | 0.060 | 0.000 | 0.234 | 0.234 | 0.386 | -1.934 | 0.518 |
| | SN | 0.139 | 0.123 | 0.055 | 0.100 | 0.000 | 0.334 | 0.334 | 0.498 | -1.419 | |
| **Trichodina sp. abundance** | HB | 19.884 | 21.188 | 8.650 | 14.460 | 1.530 | 59.180 | 57.650 | 0.877 | -0.873 | 0.792 |
| | SN | 25.904 | 33.968 | 15.191 | 10.251 | 0.230 | 80.300 | 80.070 | 0.646 | -1.551 | |
| **Glugea anomala abundance** | HB | 0.102 | 0.166 | 0.068 | 0.020 | 0.000 | 0.420 | 0.420 | 1.048 | -0.695 | 0.690 |
| | SN | 0.046 | 0.064 | 0.029 | 0.000 | 0.000 | 0.130 | 0.130 | 0.352 | -2.138 | |
| **Relative roughness** | HB | 0.003 | 0.004 | 0.001 | 0.002 | 0.001 | 0.011 | 0.009 | 1.345 | -0.108 | 0.247 |
| | SN | 0.006 | 0.006 | 0.003 | 0.006 | 0.002 | 0.016 | 0.014 | 0.717 | -1.326 | |

**Tabel S8** Explanation power of sampling year (2017) discharge and three years average (2015-2017) discharge to population genetic variance in the redundancy analysis

| Year(s) | Adjusted $R^2$ of averaged discharge | Adjusted $R^2$ of discharge variance |
|---|---|---|
| 2015-2017 | 0.0237 | 0.0218 |
| 2017 | 0.0243 | 0.0248 |

**Table S9** Correlation between 19 environmental variables measured along all sampling sites. * Significant at P < 0.05; ** Significant at P < 0.01

| EnvVar | Bankfull channel width | Bankfull channel depth | Sinuosity | Number of barriers downstream | Downstream river distance | Upstream river distance | Average conductivity | Average COD | Average pH | Average oxygen concentration | Average temperature | Bed topographical variance | Annual variance of daily discharge | Annual averaged daily discharge | Annual averaged unit discharge | Gyrodactylus sp. abundance | Trichodina sp. abundance | Glugea anomala abundance | Relative roughness |
|---|---|---|---|---|---|---|---|---|---|---|---|---|---|---|---|---|---|---|---|
| Bankfull channel width | NA | 0.27** | -0.15* | 0.09 | -0.38** | 0.53** | 0.54** | 0.18* | -0.22** | -0.36** | -0.59** | 0.18* | 0.11 | 0.07 | -0.45** | 0.32** | 0.38** | 0.25** | 0.16* |
| Bankfull channel depth | 0.27** | NA | 0.48** | -0.02 | -0.28** | -0.14 | 0.06 | 0 | -0.1 | -0.23** | -0.38** | 0.85** | -0.32** | -0.28** | -0.38** | 0.07 | -0.47** | -0.01 | 0.79** |
| Sinuosity | -0.15* | 0.48** | NA | -0.02 | 0.04 | 0.14 | 0.25** | -0.30** | 0.31** | 0.33** | -0.11 | 0.51** | -0.23** | 0.05 | 0.13 | 0.05 | -0.46** | 0.62** | 0.55** |
| Number of barriers downstream | 0.09 | -0.02 | -0.02 | NA | 0.81** | -0.28** | 0.51** | -0.53** | 0.27** | 0.66** | -0.18* | 0.09 | -0.35** | -0.24** | -0.24** | -0.18* | -0.09 | 0.34** | 0.09 |
| Downstream river distance | -0.38** | -0.28** | 0.04 | 0.81** | NA | -0.37** | 0.20** | -0.59** | 0.31** | 0.76** | 0.1 | -0.04 | -0.30** | -0.18* | 0.05 | -0.13 | -0.14 | 0.25** | 0 |
| Upstream river distance | 0.53** | -0.14 | 0.14 | -0.28** | -0.37** | NA | 0.35** | 0.19* | -0.06 | -0.43** | -0.44** | -0.18* | 0.28** | 0.30** | 0.06 | 0.12 | 0.50** | 0.32** | -0.17* |
| Average conductivity | 0.54** | 0.06 | 0.25** | 0.51** | 0.20** | 0.35** | NA | -0.12 | 0.61** | 0.36** | -0.15* | 0.04 | -0.45** | -0.29** | -0.53** | 0.1 | 0.35** | 0.81** | 0.05 |
| Average COD | 0.18* | 0 | -0.30** | -0.53** | -0.59** | 0.19* | -0.12 | NA | -0.23** | -0.59** | -0.05 | -0.09 | 0.04 | -0.1 | -0.30** | 0.18* | 0.08 | -0.32** | -0.11 |
| Average pH | -0.22** | -0.1 | 0.31** | 0.27** | 0.31** | -0.06 | 0.61** | -0.23** | NA | 0.54** | 0.58** | -0.19* | -0.73** | -0.59** | -0.37** | -0.14 | 0.23** | 0.58** | -0.18* |
| Average oxygen concentration | -0.36** | -0.23** | 0.33** | 0.66** | 0.76** | -0.43** | 0.36** | -0.59** | 0.54** | NA | 0.38** | -0.07 | -0.37** | -0.15* | 0.04 | 0.02 | -0.27** | 0.58** | 0.01 |
| Average temperature | -0.59** | -0.38** | -0.11 | -0.18* | 0.1 | -0.44** | -0.15* | -0.05 | 0.58** | 0.38** | NA | -0.48** | -0.41** | -0.44** | -0.09 | 0.05 | 0.07 | -0.04 | -0.45** |

| | | | | | | | | | | | | | | | | | | | |
|---|---|---|---|---|---|---|---|---|---|---|---|---|---|---|---|---|---|---|---|
| Bed topographical variance | 0.18* | 0.85** | 0.51** | 0.09 | -0.04 | -0.18* | 0.04 | -0.09 | -0.19* | -0.07 | -0.48** | NA | -0.19* | -0.09 | -0.20** | -0.03 | -0.40** | 0.12 | 0.99** |
| Annual variance of daily discharge | 0.11 | -0.32** | -0.23** | -0.35** | -0.30** | 0.28** | -0.45** | 0.04 | -0.73** | -0.37** | -0.41** | -0.19* | NA | 0.95** | 0.78** | -0.08 | 0.09 | -0.26** | -0.19* |
| Annual averaged daily discharge | 0.07 | -0.28** | 0.05 | -0.24** | -0.18* | 0.30** | -0.29** | -0.1 | -0.59** | -0.15* | -0.44** | -0.09 | 0.95** | NA | 0.84** | -0.07 | -0.03 | 0.02 | -0.07 |
| Annual averaged unit discharge | -0.45** | -0.38** | 0.13 | -0.24** | 0.05 | 0.06 | -0.53** | -0.30** | -0.37** | 0.04 | -0.09 | -0.20** | 0.78** | 0.84** | NA | -0.29** | -0.16* | -0.13 | -0.18* |
| Gyrodactylus sp. abundance | 0.32** | 0.07 | 0.05 | -0.18* | -0.13 | 0.12 | 0.1 | 0.18* | -0.14 | 0.02 | 0.05 | -0.03 | -0.08 | -0.07 | -0.29** | NA | -0.16* | 0.16* | 0.06 |
| Trichodina sp. abundance | 0.38** | -0.47** | -0.46** | -0.09 | -0.14 | 0.50** | 0.35** | 0.08 | 0.23** | -0.27** | 0.07 | -0.40** | 0.09 | -0.03 | -0.16* | -0.16* | NA | 0.13 | -0.41** |
| Glugea anomala abundance | 0.25** | -0.01 | 0.62** | 0.34** | 0.25** | 0.32** | 0.81** | -0.32** | 0.58** | 0.58** | -0.04 | 0.12 | -0.26** | 0.02 | -0.13 | 0.16* | 0.13 | NA | 0.19** |
| Relative roughness | 0.16* | 0.79** | 0.55** | 0.09 | 0 | -0.17* | 0.05 | -0.11 | -0.18* | 0.01 | -0.45** | 0.99** | -0.19* | -0.07 | -0.18* | 0.06 | -0.41** | 0.19** | NA |

**Table S10** Variables were selected by stepwise RDA model and their VIF value, model adjusted R square and REPI. The variable with the highest VIF value is bold-highlighted in the table. Model 2 removed the highest VIF variable from Model 1, and Model 3 removed the highest VIF variable in Model 2. In each model, from left to right, variables are ranked according to their contribution to the RDA model. * partial RDA model by using downstream distance as control variable.

| Model | Variables from left to right ranked from high to low contribution to the RDA model | | | | | | | | | Adjusted R square | REPI |
|---|---|---|---|---|---|---|---|---|---|---|---|
| 1 | **Annual variance of daily discharge** | Number of barriers downstream | Annual averaged unit discharge | Average pH | Bed topographical variance | *Glugea anomala* abundance | Average oxygen concentration | Average COD | Downstream river distance | 0.131596 | 0.726708 |
| | *20.989867* | 6.473695 | 11.362508 | 9.795661 | 2.764642 | 4.319331 | 5.891915 | 2.429945 | 6.806531 | | |
| 2 | Annual averaged unit discharge | **Downstream river distance** | Average pH | Bed topographical variance | *Glugea anomala* abundance | Number of barriers downstream | Average COD | Average oxygen concentration | | 0.120845 | 0.667342 |
| | 2.946843 | *5.590537* | 3.153895 | 1.43728 | 2.40756 | 5.266282 | 2.427903 | 4.567509 | | 0.1015377* | 0.56072 |
| 3 | Annual averaged unit discharge | Number of barriers downstream | Average pH | Bed topographical variance | Glugea anomala abundance | Average oxygen concentration | | | | 0.111347 | 0.614888 |
| | 1.801419 | 2.297345 | 2.699479 | 1.279205 | 1.959057 | 3.568289 | | | | | |

**Table S11** Biplot scores for constraining variables of the RDA model

| Environmental Variable | RDA1 | RDA2 | RDA3 | RDA4 | RDA5 | RDA6 |
|---|---|---|---|---|---|---|
| Annual averaged unit discharge | **-0.782** | -0.0936 | -0.4081 | 0.22316 | -0.3218 | -0.215 |
| Number of barriers downstream | **0.729** | 0.20672 | -0.4768 | 0.23853 | -0.1784 | -0.3019 |
| Average pH | 0.441 | **-0.5961** | -0.2505 | -0.2294 | 0.493 | -0.2165 |
| Bed topographical variance | 0.1376 | 0.19247 | 0.5513 | 0.70849 | 0.2268 | -0.2928 |
| *Glugea anomala* abundance | 0.1822 | 0.03997 | -0.1492 | -0.2382 | 0.4572 | -0.8136 |
| Average oxygen concentration | 0.3957 | **-0.3357** | -0.3094 | -0.0241 | -0.2618 | -0.6801 |
| Average COD | -0.2596 | 0.29216 | 0.4057 | -0.2959 | 0.2093 | 0.5522 |
| Eigenvalue (constrained) | 2456.36 | 1771.87 | 1552.91 | 1319.24 | 1194.82 | 918.239 |
| ProportionExplained | 0.2428 | 0.1752 | 0.1535 | 0.1304 | 0.1181 | 0.09077 |
| CumulativeProportion | 0.2428 | 0.418 | 0.5715 | 0.7019 | 0.82 | 0.91078 |

**Table S12** Top 10 outlier loci on axis 1 of the redundancy analysis (RDA1), including chromosome, location, q-value, gene name, nearest distance to SNP and function.

| No. | Chr | Location | q-value | Gene (Ref: GCF_016920845.1) | Nearest distance to SNP (bp) | Function (Description source: zfin.org) |
|---|---|---|---|---|---|---|
| 1 | chrXIII | 18916140 | 7.19397E-23 | dmrt2a | 23,933 | Predicted to have DNA-binding transcription factor activity. Involved in apoptotic process; determination of left/right symmetry; and skeletal muscle fiber development. Localizes to nucleus. Is expressed in several structures, including axis; mesoderm; mucus secreting cell; muscle; and tail bud. Orthologous to human DMRT2 (doublesex and mab-3 related transcription factor 2). |
| | | | | dmrt3a | 33,133 | Exhibits sequence-specific DNA binding activity. Predicted to be involved in regulation of transcription, DNA-templated. Predicted to localize to nucleus. Is expressed in nervous system; neural tube; and reproductive system. Orthologous to human DMRT3 (doublesex and mab-3 related transcription factor 3). |
| | | | | smarca2 | 19,009 | Predicted to have DNA binding activity and DNA-dependent ATPase activity. Predicted to be involved in ATP-dependent chromatin remodeling and positive regulation of transcription by RNA polymerase II. Predicted to localize to nucleus. Is expressed in liver and pronephric duct. Orthologous to human SMARCA2 (SWI/SNF related, matrix associated, actin dependent regulator of chromatin, subfamily a, member 2). |

| | | | | | | |
|---|---|---|---|---|---|---|
| 2 | chrXVII | 2171360 | 5.57452E-12 | Plasma membrane calcium-transporting ATPase 1-like* | in the gene | * Here, we combined the atp2b1a and atp2b1b of zebrafish to annotate the function of this gene. Atp2b1a: Predicted to have PDZ domain binding activity and calcium transmembrane transporter activity, phosphorylative mechanism. Involved in animal organ development and sensory perception of sound. Predicted to localize to intracellular membrane-bounded organelle. Is expressed in several structures, including digestive system; hematopoietic system; midbrain neural tube; nervous system; and pleuroperitoneal region. Orthologous to human ATP2B1 (ATPase plasma membrane Ca2+ transporting 1).; Atp2b1b: Predicted to have PDZ domain binding activity and calcium transmembrane transporter activity, phosphorylative mechanism. Predicted to be involved in regulation of cytosolic calcium ion concentration. Predicted to localize to intracellular membrane-bounded organelle. Is expressed in blood; eye; liver; and pleuroperitoneal region. Orthologous to human ATP2B1 (ATPase plasma membrane Ca2+ transporting 1). |
| 3 | chrXIV | 9977703 | 6.50532E-12 | Ephrin type-A receptor 5* | in the gene | * Here, we used epha4a of zebrafish to annotate the function of this gene. Predicted to have transmembrane-ephrin receptor activity. Involved in several processes, including nervous system development; optic vesicle morphogenesis; and regulation of myelination. Predicted to localize to integral component of plasma membrane; neuron projection; and receptor complex. Is expressed in several structures, including central nervous system; germ ring; mesoderm; neural keel; and neural tube. Orthologous to human EPHA4 (EPH receptor A4). |
| 4 | chrVII | 2371176 | 2.15423E-11 | Leukotriene B4 receptor 1-like | in the gene | Exhibits leukotriene B4 receptor activity. Involved in epiboly involved in gastrulation with mouth forming second. Localizes to cell surface. Orthologous to human LTB4R (leukotriene B4 receptor). |

| | | | | | | |
|---|---|---|---|---|---|---|
| 5 | chrXII | 20623122 | 4.92613E-10 | lrif1 | 5188 | Predicted to have retinoic acid receptor binding activity. Predicted to be involved in regulation of transcription, DNA-templated. Orthologous to human LRIF1 (ligand dependent nuclear receptor interacting factor 1). |
| | | | | Ankyrin repeat, SAM and basic leucine zipper domain-containing protein 1-like | 5457 | Predicted to be involved in piRNA metabolic process. Predicted to localize to pi-body. Is expressed in oocyte stage I. Orthologous to human ASZ1 (ankyrin repeat, SAM and basic leucine zipper domain containing 1). |
| 6 | chrXVI | 17772561 | 3.3449E-09 | fer1l6 | in the gene | Involved in cardiac muscle tissue development and skeletal muscle tissue development. Predicted to localize to integral component of membrane. Is expressed in several structures, including gill; gonad; heart; integument; and muscle. Orthologous to human FER1L6 (fer-1 like family member 6). |
| 7 | chrXIII | 9758098 | 5.86833E-09 | Ceramide transfer protein-like* | in the gene | * Here, we used cert1a to annotate the function of this gene. Predicted to have ceramide 1-phosphate binding activity; ceramide 1-phosphate transfer activity; and phosphatidylinositol-4-phosphate binding activity. Predicted to be involved in ER to Golgi ceramide transport and intermembrane lipid transfer. Predicted to localize to Golgi apparatus; cytosol; and membrane. Human ortholog(s) of this gene implicated in autosomal dominant non-syndromic intellectual disability 34. Is expressed in immature eye. Orthologous to human CERT1 (ceramide transporter 1). |
| 8 | chrXIII | 14520148 | 8.18842E-09 | dock5 | in the gene | Predicted to have GTPase activator activity and guanyl-nucleotide exchange factor activity. Involved in myoblast fusion. Orthologous to human DOCK5 (dedicator of cytokinesis 5). |
| 9 | chrX | 14415051 | 3.68946E-08 | top2b | in the gene | Predicted to have DNA topoisomerase type II (double strand cut, ATP-hydrolyzing) activity. Involved in axon target recognition; retina layer formation; and retinal ganglion cell axon guidance. Predicted to localize to DNA topoisomerase type II (double strand cut, ATP-hydrolyzing) complex and nucleus. Is expressed in brain; eye; immature eye; and pharyngeal arch 3-7. |

| | | | | | | |
|---|---|---|---|---|---|---|
| | | | | | | Orthologous to human TOP2B (DNA topoisomerase II beta). |
| 10 | chrXIV | 4405571 | 3.68946E-08 | Alanine--glyoxylate aminotransferase 2, mitochondrial-like | in the gene | Predicted to have pyridoxal phosphate binding activity and transaminase activity. Is expressed in liver; notochord; pronephric duct; and pronephros. Orthologous to human AGXT2 (alanine--glyoxylate aminotransferase 2). |

**Table S13** Gene expression (E, highlighted in grey color) and related phenotypes (P, highlighted in orange color) by physiological function of the top 10 outliers of RDA1.

| No. | Cardiovascular system | | Digestive system | | Endocrine system | | Immune system | | Liver and biliary system | | Musculature system | | Nervous system | | Renal system | | Reproductive system | | Respiratory system | | Sensory system | | Visual system | | Fin | | Integument | | Neural tube | | Primary germ layer | | Somite | | Polymorphic Populations |
|---|---|---|---|---|---|---|---|---|---|---|---|---|---|---|---|---|---|---|---|---|---|---|---|---|---|---|---|---|---|---|---|---|---|---|---|
| | E | P | E | P | E | P | E | P | E | P | E | P | E | P | E | P | E | P | E | P | E | P | E | P | E | P | E | P | E | P | E | P | E | P | |
| 1 | | | | | 1 | | 1 | | 1 | | 1 | | 1 | | | | 1 | | | | | | | | | | 1 | | | | 1 | 1 | 1 | 1 | 1 |
| | | | | | 1 | | | | | | | | 1 | 1 | | | 1 | | | | 1 | | | | | 1 | | | 1 | | 1 | | | | |
| | | | 1 | | | | | | 1 | | | | | | 1 | | | | | | | | | | | | | | | | | | | | |
| 2 | | | 1 | 1 | 1 | | 1 | | 1 | | 1 | | 1 | 1 | 1 | | 1 | | 1 | 1 | 1 | 1 | 1 | | | | 1 | 1 | 1 | | 1 | | 1 | | 2 |
| 3 | | | | | | | | | | | | | 1 | 1 | | | | | | | 1 | | | | | | | | 1 | | 1 | | 1 | | 3 |
| 4 | | | | | | | | | | | | | | | | | | | | | | | | | | | | | | | | | | | 6 |
| 5 | | | | | | 1 | | | | | | | | | | | | 1 | | | | | | | | | | | | | | | | | 1 |
| | | | | | 1 | | | | | | | | | | | | 1 | | | | | | | | | | | | | | | | | | |
| 6 | 1 | | | | | | | | | | 1 | 1 | 1 | 1 | | | 1 | | 1 | | | 1 | | 1 | | 1 | 1 | | | | | | | | 3 |
| 7 | | | | | | | | | | | | 1 | | 1 | | | | | | | | 1 | | 1 | | | | | | | | | | 1 | 5 |
| 8 | | | | | | | | | | | | 1 | | | | | | | | | | | | | | | | | | | | | | 1 | 2 |
| 9 | | | 1 | | | | | | | | | | 1 | 1 | | | | | 1 | | 1 | 1 | 1 | 1 | | | | | | | | | | | 5 |
| 10 | | | 1 | | | | | | 1 | | | | | | | 1 | | | | | | | | | | | | | | | | | | | 3 |
| Sum | 1 | 0 | 4 | 1 | 4 | 1 | 2 | 0 | 4 | 0 | 3 | 3 | 6 | **6** | 2 | 1 | 5 | 1 | 3 | 1 | 4 | **4** | 2 | 3 | 0 | 2 | 3 | 1 | 3 | 0 | 4 | 1 | 3 | 3 | |

**Table S14** Top 10 outlier loci on axis 2 of the redundancy analysis (RDA2), including chromosome, location, q-value, gene name, nearest distance to SNP and function.

| No. | Chr | Location | q-value | Gene (Ref: GCF_016920845.1) | Nearest distance to the SNP (bp) | Function (Description source: zfin.org) |
|---|---|---|---|---|---|---|
| 1 | chrVIII | 11159987 | 2.98E-23 | ENSGACG00000030290 | in the gene | lncRNA |
| 2 | chrXIII | 321111 | 6.23E-20 | abca2 | in the gene | Predicted to have ATPase-coupled transmembrane transporter activity and lipid transporter activity. Predicted to be involved in lipid transport. Predicted to localize to intracellular membrane-bounded organelle. Orthologous to human ABCA2 (ATP binding cassette subfamily A member 2). |
| 3 | chrIII | 11436278 | 9.02E-20 | Leucine-rich repeat-containing protein 30-like | 3836 | Not available |
| | | | | rheb | 4897 | Predicted to have GDP binding activity; GTP binding activity; and GTPase activity. Predicted to be involved in small GTPase mediated signal transduction. Predicted to localize to plasma membrane. Orthologous to human RHEB (Ras homolog, mTORC1 binding). |
| 4 | chrXIV | 7838457 | 7.42E-18 | plgrkt | in the gene | Predicted to be involved in positive regulation of plasminogen activation. Predicted to localize to integral component of plasma membrane. Orthologous to human PLGRKT (plasminogen receptor with a C-terminal lysine). |
| 5 | chrI | 5120253 | 2.33E-17 | ENSGACG00000027020 | in the gene | lncRNA |
| 6 | chrI | 6518058 | 2.33E-17 | cGMP-dependent 3',5'-cyclic phosphodiesterase | in the gene | Not available |

| | | | | | | |
|---|---|---|---|---|---|---|
| 7 | chrIX | 16068873 | 3.92E-17 | fbxw7 | in the gene | Involved in angiogenesis; negative regulation of myelination; and regulation of signal transduction. Is expressed in glioblast (sensu Vertebrata); neural tube; neuroblast (sensu Vertebrata); spinal cord; and trunk. Orthologous to human FBXW7 (F-box and WD repeat domain containing 7). |
| 8 | chrX | 10334624 | 2.42E-15 | cd164l2 | in the gene | Predicted to localize to cytoplasmic vesicle. Is expressed in neuromast and otic sensory epithelium. Orthologous to human CD164L2 (CD164 molecule like 2). |
| 9 | chrXI | 15651563 | 9.06E-15 | Protein kinase C beta type-like* | in the gene | Here, we included both prkcba and prkcbb genes to annotate the function of this gene. prkcba: Predicted to have protein serine/threonine kinase activity. Predicted to be involved in intracellular signal transduction and peptidyl-serine phosphorylation. Human ortholog(s) of this gene implicated in dilated cardiomyopathy and lung non-small cell carcinoma. Is expressed in several structures, including brain; forebrain neural keel; otic placode; pectoral fin musculature; and pronephros. Orthologous to human PRKCB (protein kinase C beta). prkcbb: Predicted to have several functions, including androgen receptor binding activity; histone kinase activity (H3-T6 specific); and nuclear receptor transcription coactivator activity. Involved in blood coagulation and embryonic hemopoiesis. Predicted to localize to nucleus. Human ortholog(s) of this gene implicated in dilated cardiomyopathy and lung non-small cell carcinoma. Is expressed in brain; retinal neural layer; and somite border. Orthologous to human PRKCB (protein kinase C beta). |
| 10 | chrIX | 19633093 | 1.65E-14 | npepps | in the gene | Exhibits aminopeptidase activity. Predicted to be involved in peptide catabolic process and proteolysis. Predicted to localize to cytoplasm. Is expressed in nervous system; pectoral fin musculature; and proliferative region. Orthologous to human NPEPPS (aminopeptidase puromycin sensitive). |

**Table S15** Gene expression (E, highlighted in grey color) and related phenotypes (P, highlighted in orange color) by physiological function of the top 10 outliers of RDA2

| No. | Cardiovascular system | | Digestive system | | Endocrine system | | Immune system | | Liver and biliary system | | Musculature system | | Nervous system | | Renal system | | Reproductive system | | Respiratory system | | Sensory system | | Visual system | | Fin | | Integument | | Neural tube | | Primary germ layer | | Somite | | Polymorphic Populations |
|---|---|---|---|---|---|---|---|---|---|---|---|---|---|---|---|---|---|---|---|---|---|---|---|---|---|---|---|---|---|---|---|---|---|---|---|
| | E | P | E | P | E | P | E | P | E | P | E | P | E | P | E | P | E | P | E | P | E | P | E | P | E | P | E | P | E | P | E | P | E | P | |
| 1 | | | | | | | | | | | | | | | | | | | | | | | | | | | | | | | | | | | 1 |
| 2 | | | | | | | | | | | | | | | | | | | | | | | | | | | | | | | | | | | 2 |
| 3 | | | | | | | | | | | | | | | | | | | | | | | | | | | | | | | | | | | 2 |
| 4 | | | | | | | | | | | | | | | | | | | | | | | | | | | | | | | | | | | 2 |
| 5 | | | | | | | | | | | | | | | | | | | | | | | | | | | | | | | | | | | 3 |
| 6 | | | | | | | | | | | | | | | | | | | | | | | | | | | | | | | | | | | 2 |
| 7 | | | | | | | | | | | | | 1 | | | | | | | | | | | | | | | | 1 | | | | | | 1 |
| 8 | | | | | | | | | | | | | 1 | | | | | | | | 1 | | | | | | | | | | | | | | 3 |
| 9 | | | 1 | | | | | | | | 1 | | 1 | 1 | 1 | | | | 1 | | 1 | 1 | 1 | 1 | 1 | | | | | | 1 | | 1 | | 9 |
| 10 | | | | | | | | | | | 1 | | 1 | | | | | | | | 1 | | 1 | | 1 | | | | | | | | | | 6 |
| Sum | 0 | 0 | 1 | 0 | 0 | 0 | 0 | 0 | 0 | 0 | 2 | 0 | **4** | 1 | 1 | 0 | 0 | 0 | 1 | 0 | **3** | 1 | 2 | 1 | 2 | 0 | 0 | 0 | 1 | 0 | 1 | 0 | 1 | 0 | |

**Table S16** Loci with strong genetic differentiation based on the habitat contrast analysis, including chromosome, locus number, gene involved, nearest distance to SNP and function.

| No. | chr | Loc | Gene | Nearest distance to the SNP (bp) | Function (Description) |
|---|---|---|---|---|---|
| 1 | chrV | 13015062 | heparan sulfate glucosamine 3-O-sulfotransferase 2-like | in the gene | Predicted to have [heparan sulfate]-glucosamine 3-sulfotransferase 1 activity. Involved in cilium assembly. Is expressed in Kupffer's vesicle; forerunner cell group; lateral plate mesoderm; nervous system; and neural tube. Orthologous to human HS3ST1 (heparan sulfate-glucosamine 3-sulfotransferase 1). |
| 2 | chrVI | 17349542 | ENSGACG00000037776 | in the gene | lncRNA |
| 3 | chrVI | 17736575 | homeobox protein Hox-D9b-like* | 3487 | * Here we used HoxD9a to annotate the gene. Predicted to have DNA-binding transcription factor activity and RNA polymerase II cis-regulatory region sequence-specific DNA binding activity. Predicted to be involved in embryonic skeletal system morphogenesis; positive regulation of transcription by RNA polymerase II; and regionalization. Predicted to localize to nucleus. Is expressed in mesenchyme; pectoral fin; pectoral fin bud; somite; and visual system. Orthologous to human HOXD9 (homeobox D9). |

| | | | | | |
|---|---|---|---|---|---|
| | | | mRNA-homeobox protein Hox-D4b-like* | 7976 | * Here we used HoxD4a to annotate the function of this gene. Predicted to have DNA-binding transcription activator activity, RNA polymerase II-specific; RNA polymerase II regulatory region sequence-specific DNA binding activity; and activating transcription factor binding activity. Involved in blood vessel morphogenesis and hemopoiesis. Predicted to localize to nucleus. Is expressed in several structures, including nervous system; neural keel; neural plate; presumptive structure; and vasculature. Orthologous to human HOXD4 (homeobox D4). |
| 4 | chrVII | 28561399 | ENSGACG00000026032 | in the gene | lncRNA |
| 5 | chrVIII | 20019021 | GTP-binding protein REM 2-like | in the gene | Exhibits calcium channel regulator activity. Involved in dopaminergic neuron differentiation. Predicted to localize to plasma membrane. Is expressed in eye and optic tectum. Orthologous to human REM2 (RRAD and GEM like GTPase 2). |
| 6 | chrX | 14811686 | ENSGACG00000033731 | in the gene | lncRNA |
| 7 | chrXI | 10192766 | uncharacterized LOC120827769 | in the gene | uncharacterized protein coding gene |
| 8 | chrXV | 3568511 | arhgef10 | in the gene | Predicted to have Rho guanyl-nucleotide exchange factor activity. Predicted to be involved in actin cytoskeleton organization and positive regulation of stress fiber assembly. Predicted to localize to centrosome and cytosol. Orthologous to human ARHGEF10 (Rho guanine nucleotide exchange factor 10). |

| | | | | | |
|---|---|---|---|---|---|
| 9 | chrXVII | 7411623 | wnk2 | in the gene | Predicted to have chloride channel inhibitor activity; potassium channel inhibitor activity; and protein serine/threonine kinase activity. Predicted to be involved in several processes, including intracellular signal transduction; protein phosphorylation; and regulation of metal ion transport. Predicted to localize to cytosol. Orthologous to human WNK2 (WNK lysine deficient protein kinase 2). |
| 10 | chrXVII | 15837830 | potassium channel subfamily K member 15-like | in the gene | Predicted to have potassium ion leak channel activity. Predicted to be involved in potassium ion transmembrane transport and stabilization of membrane potential. Predicted to localize to integral component of plasma membrane. Orthologous to human KCNK15 (potassium two pore domain channel subfamily K member 15). |
| 11 | chrXXI | 13675714 | adcyap1a | 5293 | Predicted to have neuropeptide hormone activity; peptide hormone receptor binding activity; and pituitary adenylate cyclase activating polypeptide activity. Involved in brain development and camera-type eye development. Predicted to localize to neuron projection and perikaryon. Is expressed in several structures, including central nervous system; germ ring; hatching gland; head; and optic vesicle. Orthologous to human ADCYAP1 (adenylate cyclase-activating polypeptide 1). |
| | | | mettl4 | 34586 | Predicted to have methyltransferase activity and nucleic acid binding activity. Predicted to be involved in methylation. Orthologous to human METTL4 (methyltransferase like 4). |
| 12 | chrXXI | 14600788 | ENSGACG00000004870 | in the gene | uncharacterized protein coding gene |

**Table S17** Genetic expression (E, highlighted in grey color) and related phenotypes (P, highlighted in orange color) by physiological functions of the genes listed in the habitat contrast analysis (Table S16).

| No. | Cardiovascular system | | Digestive system | | Endocrine system | | Immune system | | Liver and biliary system | | Musculature system | | Nervous system | | Renal system | | Reproductive system | | Respiratory system | | Sensory system | | Visual system | | Fin | | Integument | | Neural tube | | Primary germ layer | | Somite | |
|---|---|---|---|---|---|---|---|---|---|---|---|---|---|---|---|---|---|---|---|---|---|---|---|---|---|---|---|---|---|---|---|---|---|---|
| | E | P | E | P | E | P | E | P | E | P | E | P | E | P | E | P | E | P | E | P | E | P | E | P | E | P | E | P | E | P | E | P | E | P |
| 1 | | | | 1 | | | | | | | | | 1 | | | | | | | | 1 | | | | | | | | 1 | | 1 | | | |
| 2 | | | | | | | | | | | | | | | | | | | | | | | | | | | | | | | | | | |
| 3 | | | | | 1 | | | | | | | | 1 | | | | | | | | 1 | | 1 | | 1 | | | | | | | | 1 | |
| | 1 | 1 | 1 | | | | | | | | | | 1 | | | | | | 1 | | 1 | | | | 1 | | | | 1 | | 1 | 1 | | |
| 4 | | | | | | | | | | | | | | | | | | | | | | | | | | | | | | | | | | |
| 5 | | | | | | | | | | | | | 1 | 1 | | | | | | | 1 | | 1 | | | | | | | | | | | |
| 6 | | | | | | | | | | | | | | | | | | | | | | | | | | | | | | | | | | |
| 7 | | | | | | | | | | | | | | | | | | | | | | | | | | | | | | | | | | |
| 8 | | | | | | | | | | | | | | | | | | | | | | | | | | | | | | | | | | |
| 9 | | | | | | | | | | | | | | | | | | | | | | | | | | | | | | | | | | |
| 10 | | | | | | | | | | | | | | | | | | | | | | | | | | | | | | | | | | |
| 11 | | | 1 | | 1 | | | | | | | | 1 | 1 | | | 1 | | | | 1 | 1 | 1 | 1 | | | | | 1 | | 1 | | | |
| 12 | | | | | | | | | | | | | | | | | | | | | | | | | | | | | | | | | | |
| Sum | 1 | 1 | 2 | 1 | 2 | 0 | 0 | 0 | 0 | 0 | 0 | 0 | 5 | 2 | 0 | 0 | 1 | 0 | 1 | 0 | 5 | 1 | 3 | 1 | 2 | 0 | 0 | 0 | 3 | 0 | 3 | 1 | 1 | 0 |

### Figures

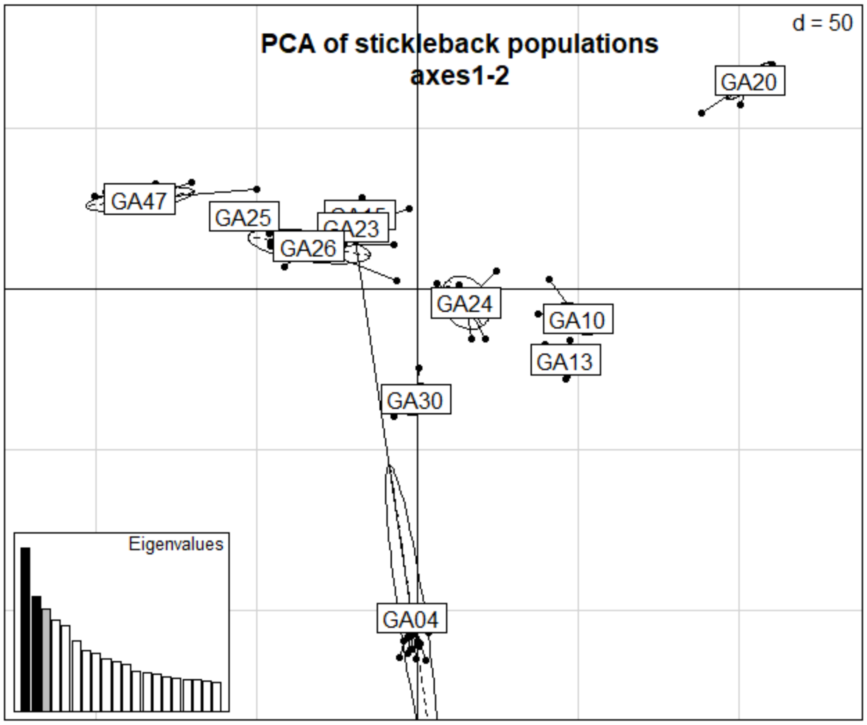

**Figure S1** PCA plot with the mislabeled individual. For sample site abbreviations see Table S1.

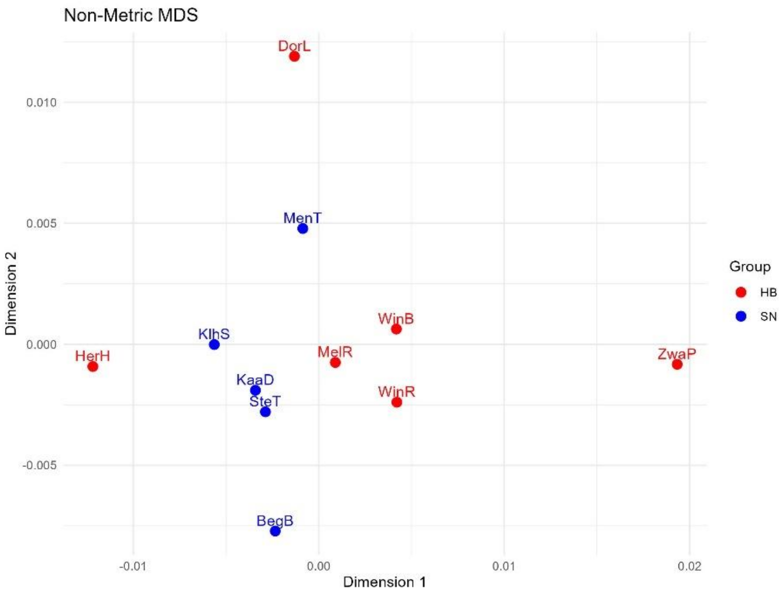

**Figure S2** Population structure calculated by non-metric multidimensional scaling.

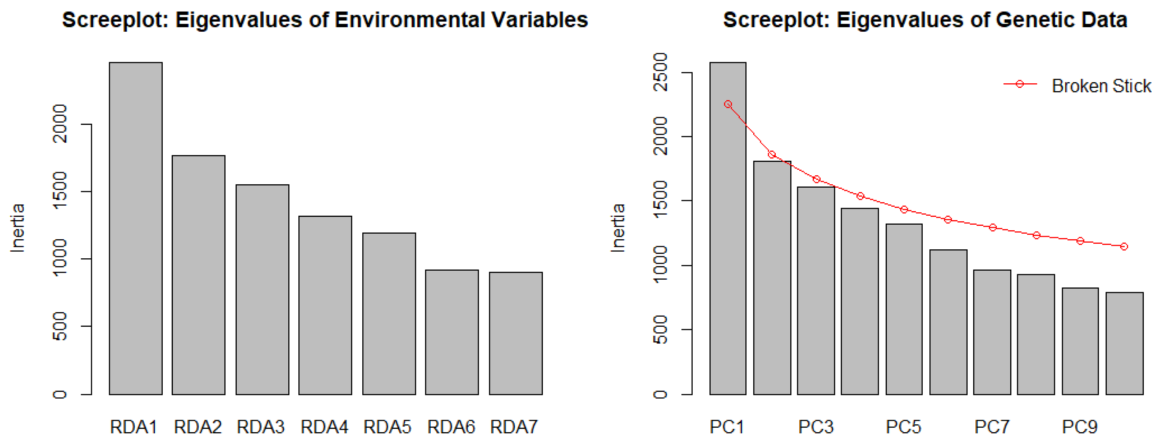

**Figure S3** Screeplot of environmental variables and genetic data of the LFMM analysis.

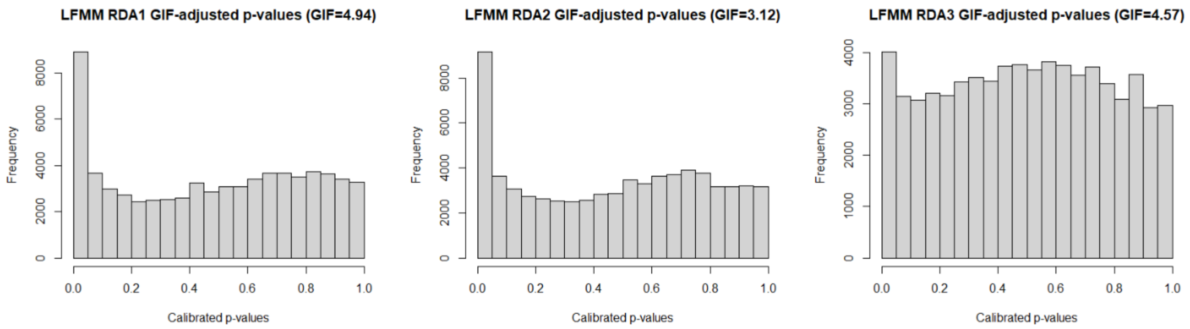

**Figure S4** GIF-adjusted p values of RDA axis 1 to 3.

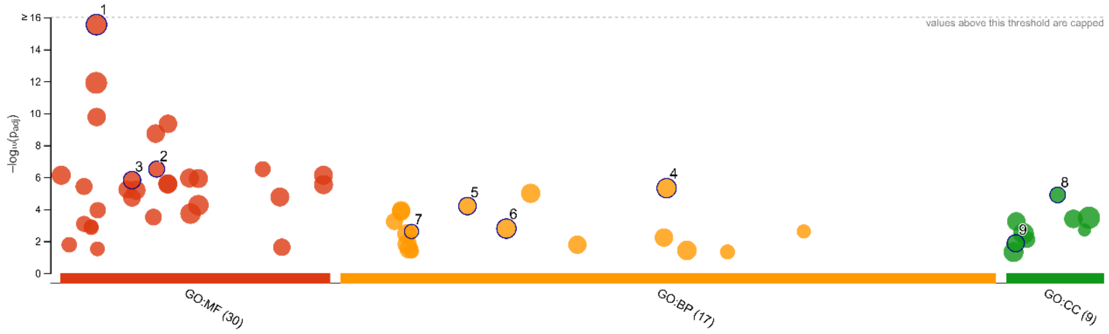

| ID | Source | Term ID | Term Name | $p_{adj}$ (query_1) |
|---|---|---|---|---|
| 1 | GO:MF | GO:0005515 | protein binding | $2.846\times10^{-16}$ |
| 2 | GO:MF | GO:0030695 | GTPase regulator activity | $3.101\times10^{-7}$ |
| 3 | GO:MF | GO:0016773 | phosphotransferase activity, alcohol group as acce... | $1.506\times10^{-6}$ |
| 4 | GO:BP | GO:0048856 | anatomical structure development | $4.797\times10^{-6}$ |
| 5 | GO:BP | GO:0016310 | phosphorylation | $6.330\times10^{-5}$ |
| 6 | GO:BP | GO:0023052 | signaling | $1.626\times10^{-3}$ |
| 7 | GO:BP | GO:0007411 | axon guidance | $2.447\times10^{-3}$ |
| 8 | GO:CC | GO:0045202 | synapse | $1.277\times10^{-5}$ |
| 9 | GO:CC | GO:0005856 | cytoskeleton | $1.342\times10^{-2}$ |

**version** e111_eg58_p18_f463989d
**date** 1/10/2025, 10:29:51 AM
**organism** gaculeatus

g:Profiler

**Figure S5** Enrichment GO plot with all outliers of molecular function (MF), biological process (BP) and cellular component (CC) are drawn in a circle, and driven terms are highlighted with a black border.

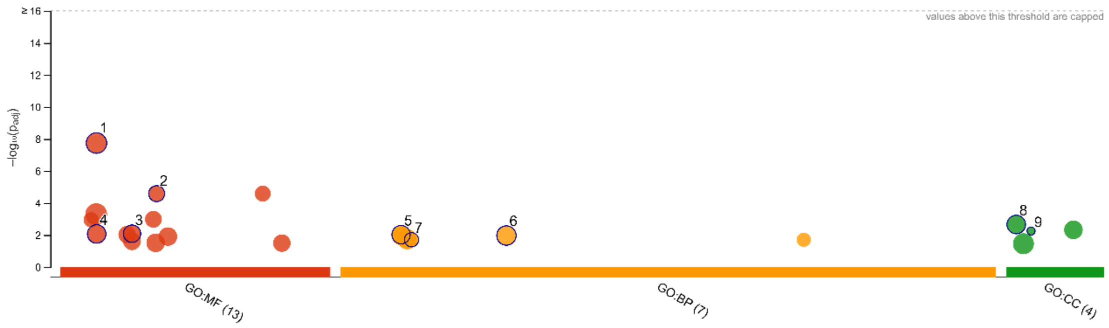

| ID | Source | Term ID | Term Name | $p_{adj}$ (query_1) |
|---|---|---|---|---|
| 1 | GO:MF | GO:0005515 | protein binding | $1.793\times10^{-8}$ |
| 2 | GO:MF | GO:0030695 | GTPase regulator activity | $2.583\times10^{-5}$ |
| 3 | GO:MF | GO:0016773 | phosphotransferase activity, alcohol group as acce... | $8.195\times10^{-3}$ |
| 4 | GO:MF | GO:0005524 | ATP binding | $8.575\times10^{-3}$ |
| 5 | GO:BP | GO:0006793 | phosphorus metabolic process | $9.435\times10^{-3}$ |
| 6 | GO:BP | GO:0023052 | signaling | $1.071\times10^{-2}$ |
| 7 | GO:BP | GO:0007411 | axon guidance | $1.959\times10^{-2}$ |
| 8 | GO:CC | GO:0005886 | plasma membrane | $2.194\times10^{-3}$ |
| 9 | GO:CC | GO:0031080 | nuclear pore outer ring | $5.574\times10^{-3}$ |

| | |
|---|---|
| **version** | e111_eg58_p18_f463989d |
| **date** | 1/10/2025, 10:38:16 AM |
| **organism** | gaculeatus |

g:Profiler

**Figure S6** Enrichment GO plot with RDA1 outliers of molecular function (MF), biological process (BP) and cellular component (CC) are drawn in a circle, and driven terms are highlighted with a black border.

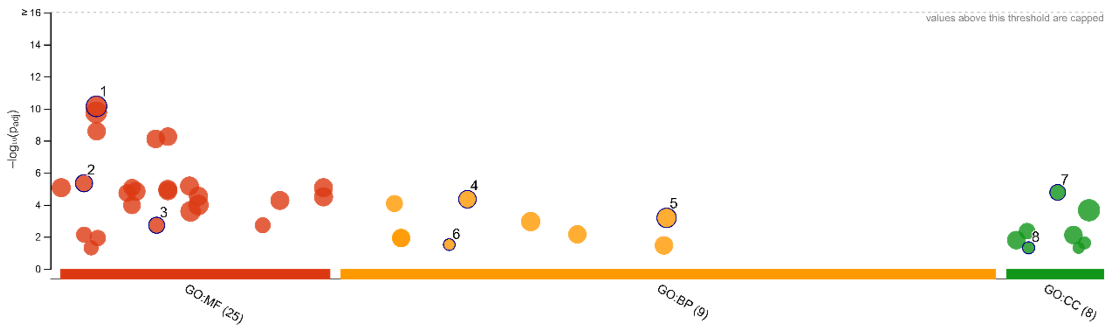

| ID | Source | Term ID | Term Name | $p_{adj}$ (query_1) |
|---|---|---|---|---|
| 1 | GO:MF | GO:0005515 | protein binding | $7.261 \times 10^{-11}$ |
| 2 | GO:MF | GO:0004672 | protein kinase activity | $4.589 \times 10^{-6}$ |
| 3 | GO:MF | GO:0030695 | GTPase regulator activity | $1.916 \times 10^{-3}$ |
| 4 | GO:BP | GO:0016310 | phosphorylation | $4.564 \times 10^{-5}$ |
| 5 | GO:BP | GO:0048856 | anatomical structure development | $6.640 \times 10^{-4}$ |
| 6 | GO:BP | GO:0010842 | retina layer formation | $3.115 \times 10^{-2}$ |
| 7 | GO:CC | GO:0045202 | synapse | $1.702 \times 10^{-5}$ |
| 8 | GO:CC | GO:0030425 | dendrite | $4.924 \times 10^{-2}$ |

| | |
|---|---|
| **version** | e111_eg58_p18_f463989d |
| **date** | 1/10/2025, 10:41:28 AM |
| **organism** | gaculeatus |

g:Profiler

**Figure S7** Enrichment GO plot with RDA2 outliers of molecular function (MF), biological process (BP) and cellular component (CC) are drawn in a circle, and driven terms are highlighted with a black border.

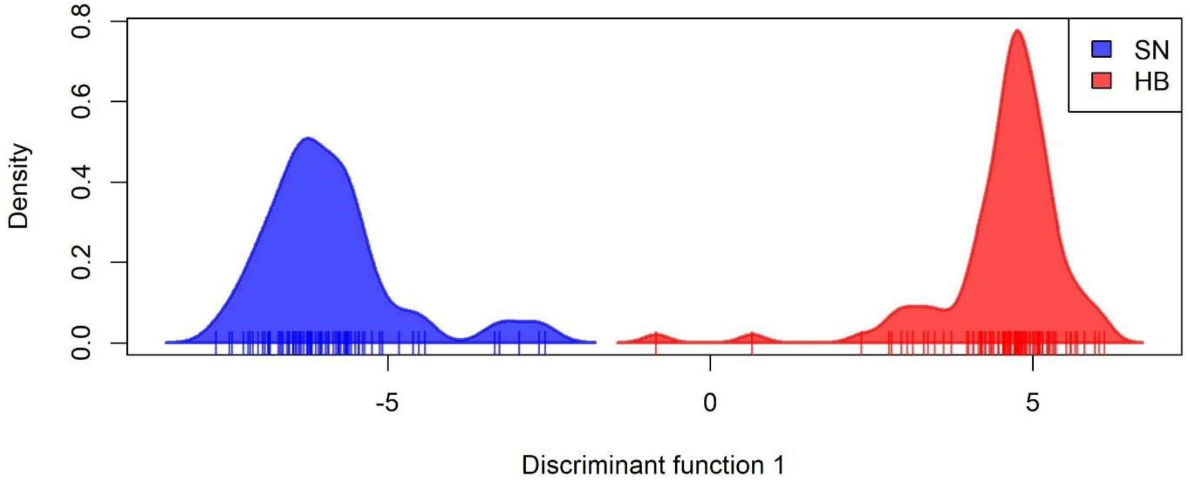

**Figure S8** Scatter plot of the DAPC of the two habitats (see legend)
